# A yeast platform for high-level synthesis of natural and unnatural tetrahydroisoquinoline alkaloids

**DOI:** 10.1101/863506

**Authors:** Michael E. Pyne, Kaspar Kevvai, Parbir S. Grewal, Lauren Narcross, Brian Choi, Leanne Bourgeois, John E. Dueber, Vincent J. J. Martin

## Abstract

The tetrahydroisoquinoline (THIQ) moiety is a privileged substructure of many bioactive natural products and semi-synthetic analogues. The plant kingdom manufactures more than 3,000 THIQ alkaloids, including the opioids morphine and codeine. While microbial species have been engineered to synthesize a few compounds from the benzylisoquinoline alkaloid (BIA) family of THIQs, low product titers impede industrial viability and limit access to the full chemical space. Here we report a THIQ platform by increasing yeast production of the central BIA intermediate (*S*)-reticuline to more than 3 g L^-1^, a 38,000-fold improvement over our first-generation strain. Gains in BIA output coincided with the formation of several substituted THIQs derived from host amino acid catabolism. Enabled by this activity, we repurposed the yeast Ehrlich pathway and demonstrate the synthesis of an array of unnatural THIQ scaffolds. This work provides a blueprint for synthesizing new privileged structures and will enable the targeted overproduction of thousands of THIQ products, including natural and semi-synthetic opioids.

---

Specialized metabolites equip plants with a chemical framework for communication and defense, whilst many such natural products have been exploited for use as flavors, dyes, and pharmaceuticals^1^. Crop-based manufacturing has enabled production of some plant metabolites at commercial scale^2^, yet low yields limit access to many high-value products. Although some natural products can be attained through chemical synthesis, the structural complexity of plant metabolites and a lack of stereocontrol result in unfeasible synthetic routes. Microbial biosynthesis has potential to overcome many of these hurdles and holds promise as a viable alternative to traditional modes of chemical and pharmaceutical manufacturing. Landmark successes in this arena, such as industrial-scale production of the anti-malarial precursor artemisinic acid^3^ and the chemical building blocks 1,3-propanediol^4^ and β-farnesene^5^, have paved the way for new opportunities in microbial biomanufacturing.

The tetrahydroisoquinoline (THIQ) structural moiety forms the basis of more than 3,000 plant natural products^6^, as well as a suite of synthetic and semi-synthetic pharmaceuticals. Naturally-occurring THIQ metabolites include the benzylisoquinoline, phenethylisoquinoline, ipecac, and Amaryllidaceae alkaloid classes^6^. The peyote cactus produces a number of simple substituted and unsubstituted THIQs^7^, while species of *Erythrina* synthesize complex spiro-THIQ alkaloids^8^. Each of these metabolite classes possesses the privileged THIQ substructure that imparts a vast array of bioactivities following derivatization in downstream tailoring reactions. The Amaryllidaceae alkaloid galantamine and the phenethylisoquinoline colchicine are commercial THIQ-derived drugs used in the treatment of Alzheimer’s disease and gout, respectively^6^. Synthetic and semi-synthetic THIQs approved by the FDA include the anti-Parkinson drug apomorphine, the chemotherapeutic trabectedin, the muscle relaxant cisatracurium, the anti-parasitic praziquantel, and the anti-chorea tetrabenazine used in the treatment of Huntington’s disease^9^.

The benzylisoquinoline alkaloids (BIAs) are the largest class of THIQ natural products and include several of the most important human medicines^10^. Morphine, codeine, and their analogues are potent BIA analgesics included in the World Health Organization’s List of Essential Medicines^11^. Papaverine is a vasodilator and antispasmodic drug, and noscapine exhibits promising anticancer properties^10^. Although global demand for morphinan BIAs is presently met through extraction from opium poppy (*Papaver somniferum*), most BIAs do not accumulate to sufficient concentrations in plant tissues. To begin exploring this untapped natural diversity, plant pathways mediating synthesis of noscapine, sanguinarine, morphine, codeine, and hydrocodone have been reconstructed in yeast^12–15^. Despite these achievements, present yeast BIA titers have been limited to less than 2 mg L^-1^ (ref. ^16–19^). For instance, the hydrocodone pathway has been reconstituted within a single yeast strain at a titer of only 0.0003 mg L^-1^ (ref. ^17^) largely due to inefficiencies in formation of the dedicated THIQ precursor. Higher levels of the key intermediate (*S*)-reticuline have been reached in *E. coli* cultures (160 mg L^-1^)^20^, yet bacteria lack membrane-bound organelles required for functional expression of the numerous cytochrome P450 enzymes in downstream BIA pathways. Consequently, engineering *E. coli* for total biosynthesis of morphinan BIAs required partitioning the pathway amongst four engineered strains^21^. Scalable production of BIAs would be facilitated through the reconstruction of heterologous pathways in a single yeast strain.

An emerging objective of synthetic biology is directed at expanding natural product diversity by engineering structural scaffolds and chemical modifications that are not observed in nature^22^. Although many biosynthetic enzymes exhibit broad substrate specificities when assayed *in vitro*, natural pathways have evolved a preference for a single substrate or very small subset of accepted building blocks^23^. The Pictet-Spengler condensation between an aryl amine and a carbonyl compound underlies the synthesis of more than 3,000 THIQ alkaloids^6^, yet much of this diversity arises from only four aldehyde species (4-hydroxyphenylacetaldehyde, 4-hydroxydihydrocinnamaldehyde, protocatechuic aldehyde, and secologanin). Because Pictet-Spenglerases are able to accept a tremendous range of carbonyl substrates^8,24-27^, natural THIQ alkaloids occupy a miniscule fraction of the conceivable chemical space. To access this untapped potential, norcoclaurine synthase (NCS) has been exploited for the synthesis of novel substituted THIQs25,27, including ones derived from ketone building blocks^8^. These *in vitro* approaches involve the synthesis of complex and unusual carbonyl substrates; consequently, producing such compounds at scale presents a formidable challenge.

We set out to establish a high-level biosynthetic route to THIQ alkaloids in which diverse structures are synthesized *in vivo* from simple substrates. We increased yeast output of the committed BIA intermediate (*S*)-reticuline to 3.1 g L^-1^, exemplifying a more than 38,000-fold improvement over our previous work^16^. These improvements shorten the path to industrial-scale production of existing BIA pharmaceuticals and will enable the targeted overproduction of thousands of natural structures that remain uncharacterized from a pharmacological perspective. We further exploited the biosynthetic potential of our THIQ platform by synthesizing a suite of unnatural THIQ scaffolds from both endogenous and supplemented amino acids. These efforts expand the diversity of privileged THIQ structures beyond natural products and outline a general synthetic biology framework for building new chemical scaffolds in yeast.

## RESULTS

### A yeast platform for high-level synthesis of substituted tetrahydroisoquinolines

To improve biosynthesis of THIQs in yeast, we focused on the first committed reaction and major rate-limiting step of the canonical BIA pathway. This conversion involves condensation of 4-hydroxyphenylacetaldehyde (4-HPAA, **1**) and dopamine (**2**) by NCS, yielding (*S*)-norcoclaurine (**3**) (**Fig. 1a**). We improved production of (*S*)-norcoclaurine more than 11,000-fold compared to our previous efforts^16^ by implementing > 20 successive strain modifications to the yeast shikimate, Ehrlich, and l-tyrosine metabolic pathways (**Fig. 1b**; strain LP478). (*S*)-Norcoclaurine synthesis in our early strains was limited by 4-HPAA, which is rapidly transformed to the corresponding fusel acid (4-hydroxyphenylacetate; 4-HPAC, **4**) or alcohol (tyrosol, **5**) via the Ehrlich pathway. The specific enzymes mediating these redox reactions have thus far evaded identification^28^, as yeast produces a suite of more than 30 oxidoreductases^29^. A gene deletion screen of more than 20 of these candidates implicated the Ari1 short-chain dehydrogenase/reductase in 4-HPAA reduction (**Supplementary Fig. 1**). Owing to redundancy in yeast oxidoreductases, we proceeded to delete a total of five genes encoding NADPH-dependent reductases and dehydrogenases (in order: *ari1*Δ *adh6*Δ *ypr1*Δ *ydr541c*Δ *aad3*Δ), enabling the production of 77 mg L^-1^ of (*S*)-norcoclaurine by strain LP386 (**Fig. 1b** and **Supplementary Fig. 2**). Although the *ALD4* aldehyde-dehydrogenase-encoding gene was deleted in early strains, resulting in gains in both dopamine and (*S*)-norcoclaurine titer (**Supplementary Fig. 1** and **2**), we reintroduced the gene in later strains to promote ethanol consumption in fed-batch fermentor cultures (**Fig. 1b**; strains LP474 and LP478). To further improve precursor supply, strain LP478 incorporates overexpression of genes encoding chorismate synthase (*ARO2*), prephenate dehydrogenase (*TYR1*), and phenylpyruvate decarboxylase (*ARO10*), as well as feedback-resistant forms of 3-deoxy-d-arabino-heptulosonate-7-phosphate synthase (*ARO4^FBR^*) and chorismate mutase (*ARO7^FBR^*). We also inactivated the l-phenylalanine and l-tryptophan biosynthetic pathways through deletion of prephenate dehydratase (*pha2*Δ) and indole-3-glycerol-phosphate synthase (*trp3*Δ) genes, respectively.

**Fig. 1.**
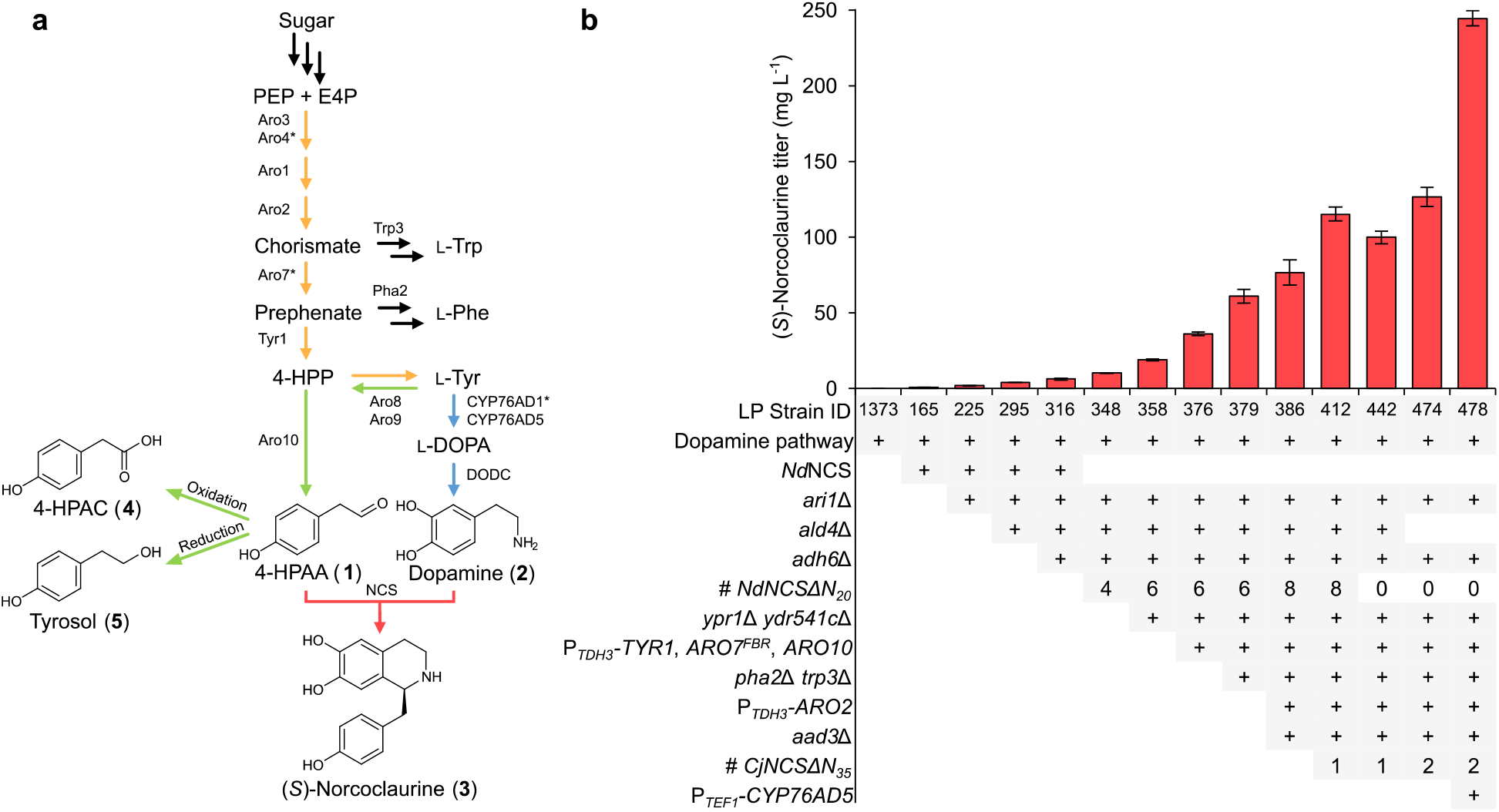
– Engineering a THIQ-producing yeast. **a,** (*S*)-Norcoclaurine (**3**) synthetic pathway in engineered yeast. The native yeast Ehrlich pathway (green) supplies 4-HPAA (**1**) from l-tyrosine, while a heterologous pathway (blue) generates dopamine (**2**), also from l-tyrosine. CYP76AD5 is a more active plant tyrosine hydroxylase compared to CYP76AD1* (CYP76AD1^W13L F309L^), which is an engineered variant of CYP76AD1. NCS catalyzes an enantioselective Pictet-Spengler condensation of 4-HPAA and dopamine, yielding (*S*)-norcoclaurine (pink). Native shikimate and l-tyrosine biosynthetic pathways are shown in orange. **b,** (*S*)-Norcoclaurine (**3**) titer in culture supernatants of successive engineered strains. Error bars represent s.d. of three biological replicates. All strains exhibited a significant increase (*P* < 0.05) in (*S*)-norcoclaurine titer relative to the respective parental strain, with the exception of strain LP442, which exhibited a significant decrease (*P* < 0.05) in titer relative to strain LP412. Refer to **Supplementary Table 4** for genotypes of the full 23-strain (*S*)-norcoclaurine lineage. Abbreviations: l-DOPA, l-3,4-dihydroxyphenylalanine; DODC, DOPA decarboxylase; E4P, erythrose-4-phosphate; 4-HPAA, 4-hydroxyphenylacetaldehyde; 4-HPAC, 4-hydroxyphenylacetate; NCS, norcoclaurine synthase; PEP, phosphoenolpyruvate; l-Phe, l-phenylalanine; l-Trp, l-tryptophan; l-Tyr, l-tyrosine.

Our initial (*S*)-norcoclaurine-producing strains contained an N-terminally truncated NCS variant from *Nandina domestica* (*Nd*NCSΔN_20_)^30^ (**Supplementary Fig. 3**). Because (*S*)-norcoclaurine production increased by supplying additional copies of the *NdNCSΔN_20_* gene (up to eight copies) (**Fig. 1b** and **Supplementary Fig. 2**), we set out to identify a more efficient NCS ortholog. Several groups have reported the use of truncation variants of NCS from *Coptis japonica* (C*j*NCS)^18,25^. We explored several extensive N-terminal truncations of C*j*NCS by reducing the protein to the core Bet v1 domain and comparing NCS activity to *Nd*NCSΔN_20_. A 35 amino acid deletion (C*j*NCSΔN_35_) facilitated the production of five-fold more (*S*)-norcoclaurine than an isogenic strain harboring *Nd*NCSΔN_20_ (**Supplementary Fig. 4**). Implementation of C*j*NCSΔN_35_ in strain LP386 possessing eight copies of *NdNCSΔN_20_* facilitated a 50% increase in (*S*)-norcoclaurine titer (strain LP412). We proceeded to delete all eight copies of *NdNCSΔN_20_* (strain LP442) and integrated an additional copy of *CjNCSΔN_35_* (strain LP474).

Concurrent precursor feeding experiments revealed a greater increase in (*S*)-norcoclaurine production upon supplementation of l-DOPA compared to l-tyrosine (**Supplementary Fig. 5**), indicating a limitation in dopamine supply. To address this substrate imbalance, we introduced a superior tyrosine hydroxylase ortholog (CYP76AD5)^31–34^ into strain LP474, which already contains an engineered CYP76AD1 variant (CYP76AD1^W13L F309L^)^16^ (**Supplementary Fig. 6**; strain LP478). In line with substrate supplementation experiments, increased expression of tyrosine hydroxylase through implementation of *CYP76AD5* doubled (*S*)-norcoclaurine titer (245 mg L^-1^) relative to strain LP474. Cultivation of strain LP478 in a pulsed sucrose fed-batch fermentor yielded 1.6 g L^-1^ of (*S*)-norcoclaurine and 2.1 g L^-1^ of tyrosol (**Supplementary Fig. 7**). Thus, applying a fed-batch process to our engineered *S. cerevisiae* strain enabled unprecedented titers of BIAs that are synthesized directly from sugar.

### Production of functionalized substituted THIQs

Having achieved > 1 g L^-1^ of (*S*)-norcoclaurine, we sought to extend the BIA pathway to the major branch point intermediate (*S*)-reticuline. This four-step conversion involves methylation of the THIQ motif of (*S*)-norcoclaurine (**3**) by two methyltransferases from opium poppy (*Papaver somniferum*; *Ps*6OMT and *Ps*CNMT), yielding (*S*)-coclaurine (**8**) followed by (*S*)-*N*-methylcoclaurine (**9**) (**Fig. 2a**). The benzyl moiety of (*S*)-*N*-methylcoclaurine is then hydroxylated by a cytochrome P450 *N*-methylcoclaurine hydroxylase from California poppy (*Eschscholzia californica*; CYP80B1 or *Ec*NMCH)^16^, yielding (*S*)-3’-hydroxy-*N*-methylcoclaurine (**10**), which is subsequently methylated by a 4’*O*-methyltransferase (*Ps*4’OMT2) to generate (*S*)-reticuline (**11**). Our (*S*)-reticuline pathway module included a cytochrome P450 reductase (CPR) from *Arabidopsis thaliana* (*At*ATR2) for electron transfer to *Ec*NMCH. Introduction of this composite pathway to strain LP478 generated 270 mg L^-1^ of (*S*)-reticuline in microtiter plate cultivations (**Fig. 2b**; strain LP490). All three intermediates in the conversion of (*S*)-norcoclaurine (**3**) to (*S*)-reticuline (**11**) were detected. Assuming an equal ionization efficiency, (*S*)-3’-hydroxy-*N*-methylcoclaurine (**10**) was found to accumulate to the greatest extent. To address this bottleneck, we integrated an additional copy of *Ps4’OMT2* (strain LP491), which nearly abolished accumulation of (*S*)-3’-hydroxy-*N*-methylcoclaurine and resulted in a 76% improvement in (*S*)-reticuline titer to 476 mg L^-1^.

**Fig. 2.**
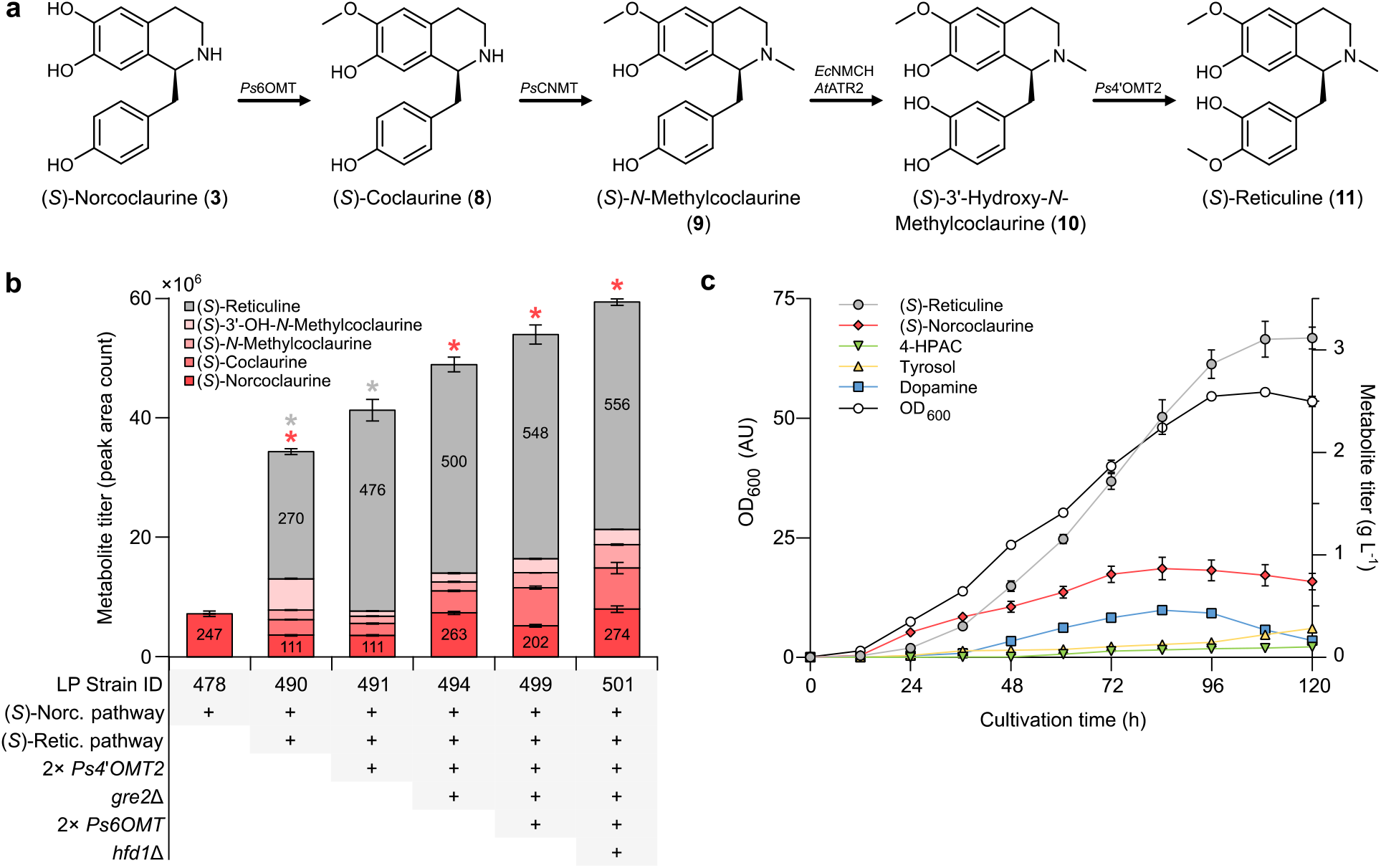
–Extending the canonical THIQ pathway to (*S*)-reticuline. **a,** Composite pathway for synthesis of (*S*)-reticuline (**11**) from (*S*)-norcoclaurine (**3**). **b,** Relative titers of (*S*)-reticuline pathway intermediates in culture supernatants of successive engineered production strains. Peak area counts are shown for all pathway intermediates and absolute titers are depicted in mg L^-1^ for (*S*)-norcoclaurine and (*S*)-reticuline. Error bars represent s.d. of three biological replicates. Asterisk (*) denotes a significant increase or decrease (*P* < 0.05) in (*S*)-norcoclaurine or (*S*)-reticuline titer relative to the parental strain. **c,** Cultivation of an (*S*)-reticuline-producing strain (LP501 harboring pHUM) in a sucrose-pulsed fed-batch fermentor. Growth of biomass (OD_600_) and accumulation of BIA metabolites in the culture medium during cultivation. Error bars represent s.e.m. of duplicate experiments.

We measured fusel products generated by strain LP491 in microtiter plate cultures and observed substantial accumulation of tyrosol (235 mg L^-1^) despite deletion of five oxidoreductase genes, implying that additional unidentified enzymes reduce 4-HPAA to tyrosol. Single-gene deletions of seven of these candidates (*AAD4, AAD14, ADH7, GCY1, GRE2, SFA1*, and *YGL039W*) in strain LP491 identified Gre2 as a potent 4-HPAA reductase (**Supplementary Fig. 8**), as its inactivation facilitated an 83% decrease in tyrosol formation in microtiter plate cultures (**Supplementary Fig. 9**; strain LP494). Although (*S*)-reticuline titer was unaltered under these conditions, (*S*)-norcoclaurine production more than doubled (111 mg L^-1^ to 263 mg L^-1^) (**Fig. 2b**). Growth of strain LP494 in a pulsed fed-batch fermentor produced only 138 mg L^-1^ of tyrosol, whereas the (*S*)-norcoclaurine-producing strain LP478 cultivated under identical conditions generated 2.1 g L^-1^ of tyrosol (**Supplementary Fig. 7**), corresponding to a 93% reduction in tyrosol synthesis with improved flux to the heterologous pathway (**Supplementary Fig. 9**).

With tyrosol production nearly abolished, 4-HPAC increased in concentration and became the dominant fusel product in both microtiter plate and pulsed fed-batch fermentor cultures (**Supplementary Fig. 9**). We thus set out to identify the enzyme(s) responsible for the oxidation of 4-HPAA to 4-HPAC. We surveyed yeast aldehyde dehydrogenases for cytosolic enzymes that exhibit activity on aromatic aldehydes. Although Ald2 and Ald3 meet these criteria, deletion of both *ALD2* and *ALD3* in strain LP492 (resulting in strain LP495) failed to alter 4-HPAC concentration in microtiter plate cultures (data not shown). We then turned our attention to Hfd1, a dual-function aldehyde dehydrogenase involved in ubiquinone (coenzyme Q10) biosynthesis and fatty acid catabolism. One of the physiological substrates of Hfd1 is 4-hydroxybenzaldehyde^35^, a close structural analogue of 4-HPAA, and localization studies have shown that Hfd1 resides in the outer mitochondrial membrane where it is ostensibly exposed to the cytosol^36^. Deletion of *HFD1* in our LP494 *gre2*Δ mutant strain (yielding strain LP498) decreased 4-HPAC production by more than 80% in both microtiter plate and pulsed fed-batch fermentor cultures (**Supplementary Fig. 10**). Our final (*S*)-reticuline production strain (LP501) incorporates *gre2*Δ and *hfd1*Δ mutations for reduced synthesis of fusel products, as well as an additional copy of *Ps6OMT* to improve conversion of (*S*)-norcoclaurine to (*S*)-coclaurine. Strain LP501 produced 3.1 g L^-1^ of (*S*)-reticuline and 0.7 g L^-1^ of (*S*)-norcoclaurine in a pulsed fed-batch fermentor (**Fig. 2c**). Tyrosol, 4-HPAC, and dopamine reached final concentrations of less than 0.3 g L^-1^.

### *De novo* synthesis of unnatural substituted tetrahydroisoquinolines

In addition to benzylisoquinolines, higher plants synthesize phenethylisoquinolines and the Amaryllidaceae alkaloids via 4-hydroxydihydrocinnamaldehyde and protocatechuic aldehyde,respectively. These natural pathways as well as *in vitro* studies^8,25,27,37^ highlight the broad substrate specificity of Pictet-Spenglerases and prompted us to employ our engineered strains to explore NCS promiscuity. We examined LC-MS spectra derived from supernatants of an (*S*)-norcoclaurine production strain for peaks indicative of substituted THIQ products. Our search consisted of 64 theoretical THIQ products derived through Pictet-Spengler condensation of dopamine and endogenous yeast carbonyl species. From this search we identified four putative LC-MS peaks (**Supplementary Table 1**). A major peak corresponding to salsolinol (**13**), derived from condensation of dopamine (**2**) and acetaldehyde (**12**) (**Fig. 3a**), was observed in supernatants of all dopamine-producing strains irrespective of the presence of an NCS biosynthetic enzyme. Three additional LC-MS peaks were consistent with substituted THIQs derived from condensation of dopamine (**2**) and aldehydes from the Ehrlich pathway^28^. These substituted THIQs (**16**, **19**, and **22**) arise from l-phenylalanine (**14**), l-tryptophan (**17**), and l-leucine (**20**) catabolism via the respective aldehydes phenylacetaldehyde (PAA, **15**), indole acetaldehyde (IAA, **18**), and 3-methylbutanal (3MB, **21**) (**Fig. 3b**). Targeted fragmentation of substituted THIQ products yielded spectra consistent with the presumed metabolite identities^38^(**Supplementary Fig. 11**). Further, cultivating strain LP385 harboring *Cj*NCSΔN_35_ on l-tyrosine (**6**), l-phenylalanine (**14**), l-tryptophan (**17**), or l-leucine (**20**) as a sole nitrogen source increased the area of LC-MS peaks corresponding to (*S*)-norcoclaurine (**3**) and products **16**, **19**, and **22**, respectively (**Supplementary Fig. 12**), providing additional evidence that these substituted THIQs arise from amino acid degradation. Analogous substituted THIQs derived from catabolism of l-isoleucine and l-valine were not observed since α-substituted aldehydes are not well tolerated by NCS^39^.

**Fig. 3.**
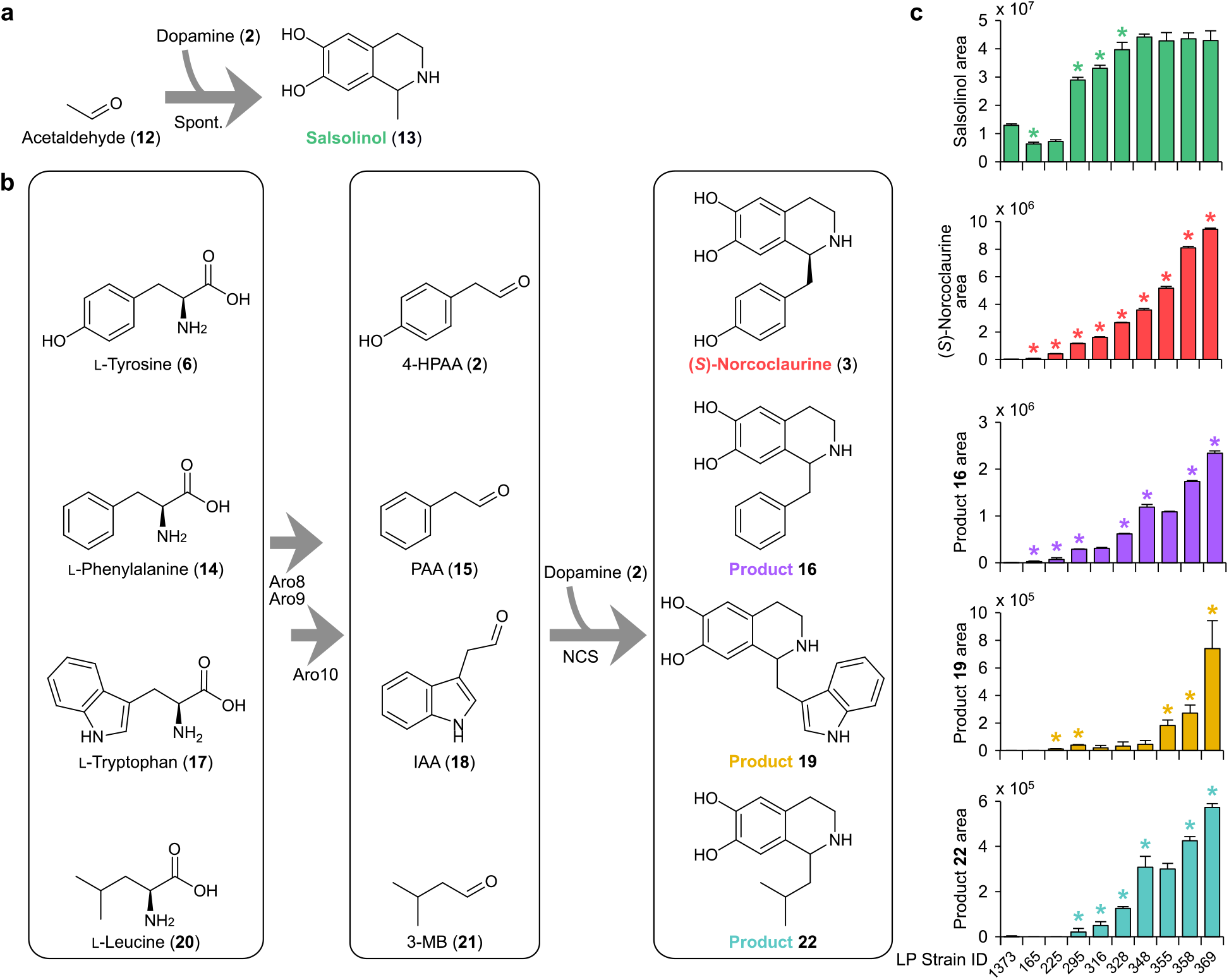
– *De novo* synthesis of substituted tetrahydroisoquinolines in strains engineered for (*S*)-norcoclaurine production. **a,** Formation of salsolinol (**13**) from acetaldehyde (**12**) and dopamine (**2**) occurs spontaneously in yeast strains engineered for dopamine production. **b,** NCS-catalyzed formation of substituted THIQs through catabolism of endogenous amino acids via the Ehrlich pathway. Amino acids are converted to the respective aldehyde species via sequential transamination (Aro8/Aro9) and decarboxylation (Aro10) reactions. In the presence of dopamine (**2**) and NCS, aldehydes are converted to the corresponding substituted THIQs. Stereochemistry of unnatural substituted THIQs is omitted. **c,** Relative levels of substituted THIQs in yeast strains engineered for (*S*)-norcoclaurine (**3**) production. Strain 1373 possesses the dopamine pathway and lacks an NCS biosynthetic enzyme. Refer to **Fig. 1b** and **Supplementary Table 4** for genotypes of engineered strains. Error bars represent s.d. of three biological replicates. Asterisk (*) denotes a significant increase or decrease (*P* < 0.05) in product titer relative to the precursor strain. Abbreviations: 4-HPAA, 4-hydroxyphenylacetaldehyde; IAA, indole acetaldehyde; 3-MB, 3-methylbutanal; PAA, phenylacetaldehyde; spont., spontaneous.

We analyzed LC-MS spectra from supernatants of our oxidoreductase gene deletion strains to monitor levels of substituted THIQ products (**16**, **19**, and **22**) relative to improvements in (*S*)-norcoclaurine concentration (**Fig. 3c**). These THIQs were not identified in supernatants of a dopamine-producing strain lacking an NCS enzyme (strain 1373) and, with the exception of **16**,were not detected in the *Nd*NCS strain used as the basis of our oxidoreductase gene deletions (strain LP165). Overall, formation of these products closely paralleled (*S*)-norcoclaurine, in which biosynthesis of **16**, **19**, and **22** increased in accordance with (*S*)-norcoclaurine throughout the oxidoreductase gene deletion lineage. Synthesis of **16** and **19** increased through deletion of *ARI1*, whereas deletion of *YPR1* or *ADH6* increased production of **19** and **22**, respectively. Deletion of *YDR541C* enhanced levels of all THIQs derived from the Ehrlich pathway.

### Synthesis of unnatural substituted tetrahydroisoquinolines from supplemented amino acids

Owing to the capacity of our engineered host to synthesize substituted THIQs from endogenous amino acids, we reasoned that supplying exogenous amino acids would enable the synthesis of additional THIQ structures. We devised a THIQ synthesis assay by cultivating our (*S*)-reticuline production strain (LP501) on individual amino acids as the major source of nitrogen. In this manner, amino acid utilization via the Ehrlich pathway directly links cell growth to aldehyde and thus THIQ formation. Using this assay, we synthesized a diverse set of THIQs possessing both aliphatic and aromatic substitutions (**Supplementary Fig. 13** and **Supplementary Table 2**). Supplying l-methionine (**23**) as the major nitrogen source generated the sulfur-containing THIQ **25** via methional (**24**) (**Fig. 4**). l-DOPA is a non-proteinogenic derivative of l-tyrosine, suggesting that it serves as a substrate for enzymes of the Ehrlich pathway. Feeding l-DOPA (**7**) to strain LP501 gave rise to norlaudanosoline (tetrahydropapaveroline, **28**) via 3,4-dihydroxyphenylacetaldehyde (3,4-dHPAA, **27**). We also demonstrated production of ethyl-, propyl-, butyl-, and pentyl-substituted THIQs (**31**, **35**, **39**, and **43**, respectively) by supplying l-2-aminobutyrate (**29**), l-norvaline (**33**), l-norleucine (**37**), and l-2-aminoheptanoic acid (**41**), respectively (**Fig. 4**). Targeted fragmentation of substituted THIQs yielded spectra consistent with the presumed metabolite identities^38^ (**Supplementary Fig. 14**).

**Fig. 4.**
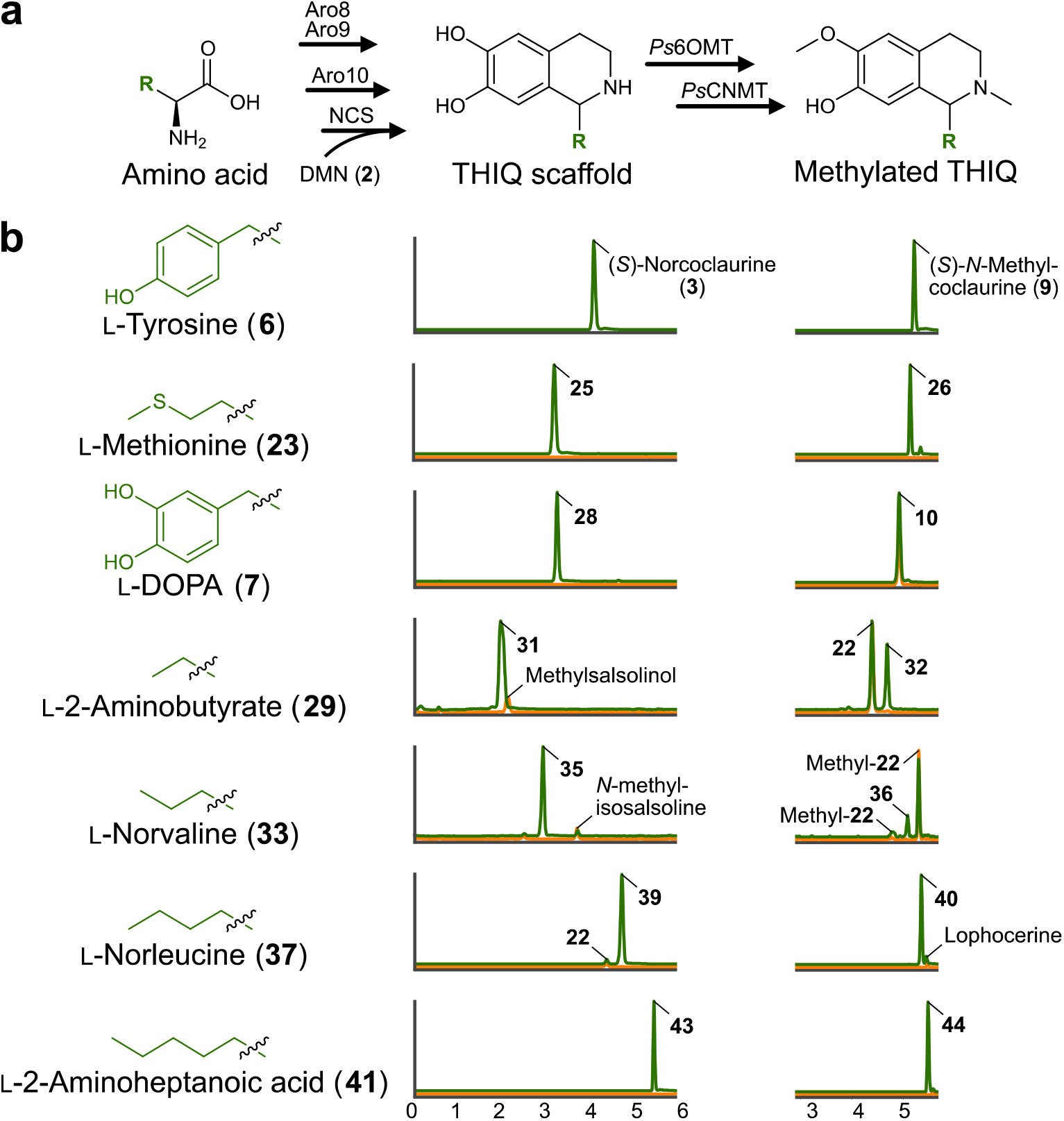
– Synthesis and modification of unnatural substituted tetrahydroisoquinolines from supplemented amino acids. **a,** NCS-catalyzed formation of substituted THIQs through catabolism of externally supplied amino acids via the Ehrlich pathway. **b,** Ion-extracted LC-MS chromatograms of strain LP501 grown on l-methionine (**23**), l-DOPA (**7**), l-2-aminobutyrate (**29**), l-norvaline (**33**), l-norleucine (**37**), or l-2-aminoheptanoic acid (**41**) as the chief source of nitrogen (green). Aldehydes derived from amino acids were incorporated into the corresponding substituted THIQs and methylated by BIA tailoring enzymes (*Ps*6OMT and *Ps*CNMT) produced by strain LP501. Substituted THIQs and their methylated derivatives shifted in retention time relative to the canonical BIA products from l-tyrosine, namely (*S*)-norcoclaurine (**3**) and (*S*)-*N*-methylcoclaurine (**9**), which were formed *de novo* on all amino acid substrates. Growth of strain LP501 on urea as the major nitrogen source (orange) failed to generate peaks corresponding to substituted THIQs with the exception of **31** and **40**, which co-elute with singly-methylated salsolinol (by *Ps*6OMT or *Ps*CNMT) and lophocerine, respectively. Product **36** elutes closely with both singly-methylated derivatives of **22** (by *Ps*6OMT or *Ps*CNMT), which are synthesized *de novo* from l-leucine. Products **35** and **39** are isomers of *N*-methylisosalsoline and **22**, respectively, but do not co-elute with these *de novo* products. Methylation of norlaudanosoline (**28**) by *Ps*6OMT and *Ps*CNMT yields 3-hydroxy-*N*-methylcoclaurine (**10**), which is produced *de novo* by strain LP501 irrespective of the nitrogen source. All *m/z* values were calculated based on the expected structures of the respective compounds of interest (**Supplementary Fig. 13** and **Supplementary Table 2**). Abbreviations: DMN, dopamine.

We also explored the promiscuity of BIA tailoring enzymes toward unnatural substituted THIQs. Enzymes involved in methylation of (*S*)-norcoclaurine (*Ps*6OMT and *Ps*CNMT) exhibited activity on all substituted THIQs derived from supplemented amino acids (**Fig. 4**), as well as THIQ scaffolds synthesized *de novo* from endogenous substrates (**Supplementary Fig. 15**). Methylation of salsolinol (**13**) by *Ps*6OMT and *Ps*CNMT yielded *N*-methylisosalsoline (**45**), while methylation of the THIQ scaffold derived from l-leucine (**22**) gave rise to lophocerine (**48**), a naturally occurring THIQ alkaloid^40^. Thus, the pool of BIAs and other THIQs that can be synthesized using our high-flux THIQ-producing yeast is immensely diverse.

## DISCUSSION

Our work demonstrates a major step towards industrial synthesis of microbially-sourced THIQ pharmaceuticals by increasing production of the key BIA intermediate (*S*)-reticuline to 3.1 g L^-1^. Many bioprocesses become commercially viable when titers reach the gram per liter level and a target of 5 g L^-1^ has been set forth for commercial-scale production of opioids^17^. Although our titers are commensurate with this benchmark, morphine is synthesized several enzymatic steps downstream of (*S*)-reticuline. The efficiency of these reactions within a high titer production host remains unknown, yet the recent discovery of dedicated thebaine and neopinone biosynthetic enzymes^41,42^ primes our engineered yeast for production of opioid pharmaceuticals at scale. Beyond natural products, our microbial production platform also enables the production of diverse molecules not sampled in nature.

Owing to their reactivity and toxicity within biological systems, microorganisms rapidly transform aldehydes to acids or alcohols^43^. Our initial BIA-producing strains were found to be limited by 4-HPAA, which is converted to tyrosol or 4-HPAC by a suite of hitherto unidentified enzymes^16,44^. We have resolved this long-standing issue by identifying six redundant oxidoreductases involved in tyrosol formation (Ari1, Adh6, Ypr1, Ydr541c, Aad3, and Gre2), in addition to the major 4-HPAC-forming enzyme (Hfd1). These oxidoreductases are involved in the catabolism of multiple Ehrlich pathway amino acids, as their inactivation triggered the biosynthesis of THIQs derived from l-phenylalanine, l-tryptophan, and l-leucine. With respect to fusel alcohol synthesis, most prior reports have implicated the ADH family of largely NADH-dependent dehydrogenases^28,45^, whilst we have shown that exclusively NADPH-dependent enzymes mediate tyrosol formation. Relatedly, identification of Hfd1 as the chief 4-HPAC-forming enzyme contrasts the widely held notion that exclusively ALD-family enzymes catalyze the oxidation of fusel aldehydes in yeast^28^. These findings have implications within the food and beverage industries, in which production of fusel products is a key determinant of flavor profiles^28^. More broadly, the capacity of our engineered strain to synthesize high levels of both aliphatic and aromatic aldehydes suggests that it could be repurposed to produce diverse compounds derived from carbonyl intermediates, for instance biofuels, fragrances, and chemotherapeutics^28,46–48^.

Our THIQ production host enables the synthesis of new privileged structures by diverting carbonyl intermediates of the yeast Ehrlich pathway to THIQ synthesis. We exploited this activity to synthesize several unnatural THIQ analogues by supplying a suite of aromatic and aliphatic building blocks to our engineered host. To our knowledge most of these structures have not been observed in nature and thus our work broadens the chemical space occupied by natural THIQ products. It is noteworthy that the isobutyl-substituted THIQ (**22**) derived from l-leucine has been identified in the senita cactus (*Lophocereus schotti*) where it forms the basis of the methylated alkaloid lophocerine^40^. Our (*S*)-reticuline production host recapitulated this natural pathway, as implementation of *Ps*6OMT and *Ps*CNMT resulted in *de novo* synthesis of lophocerine. We also observed the formation of salsolinol and its methylated derivatives, demonstrating that diverse THIQ scaffolds can be furnished with functional groups using canonical BIA tailoring enzymes. Furthermore, the THIQ scaffold formed from PAA (**16**) possesses the core BIA skeleton, indicating that the benzyl moiety of BIAs can also derive from l-phenylalanine. Products arising from **16** would lack hydroxylation at the 4’ position of the benzyl substituent that originates from 4-HPAA and is present in nearly all BIA structures characterized to date. In this context, sacred lotus (*Nelumbo nucifera*) synthesizes aporphine alkaloids possessing an unsubstituted benzyl moiety^49^ and thus our work exposes a plausible biosynthetic route to these distinct BIAs.

In summary, we engineered a yeast host capable of synthesizing high titers of the central BIA intermediate (*S*)-reticuline. This metabolite is a direct precursor to all natural BIAs, including the morphinan family, providing a platform for the microbial synthesis of natural opiates and their semi-synthetic derivatives. Microbial biosynthesis also provides the opportunity to access thousands of untapped metabolites that are present at low levels in source plants, a goal that is now within reach given our improvements in yeast THIQ biosynthesis. Finally, linking plant BIA metabolism to our repurposed Ehrlich pathway afforded yeast with the capacity to synthesize diverse new THIQ scaffolds. The THIQ motif is a privileged substructure that primes many natural products for bioactivity and thus harnessing this newfound activity expands the diversity of THIQ alkaloids beyond the canonical l-tyrosine pathway and will bring about the development of alkaloids with new pharmacological properties.

## Supporting information

Pyne et al supplemental information

## Acknowledgments

We thank Marcos DiFalco for assistance with LC-MS analyses, as well as Chris Law and the Centre for Microscopy and Cellular Imaging, which is funded by Concordia University and the Canada Foundation for Innovation. This study was financially supported by an NSERC-Industrial Biocatalysis Network (IBN) grant, an NSERC Discovery grant, and by River Stone Biotech ApS. M.E.P. was supported by an NSERC Postdoctoral Fellowship, K.K. was supported by a Concordia University Horizon Postdoctoral Fellowship, L.N. was supported by a FQRNT DE Doctoral Research Scholarship for Foreign Students, L.B. was supported by an NSER Concordia graduate scholarship. J.E.D. is supported by NSF MCB 1818307. V.J.J.M. is supported by a Concordia University Research Chair.

## Author contributions

M.E.P., K.K., P.S.G., L.N., J.E.D., and V.J.J.M. designed the research. M.E.P., K.K., P.S.G, and B.C. performed the experiments. L.N. assisted with HPLC and MS analyses and L.B. assisted in preliminary studies. V.J.J.M. and J.E.D. supervised the research. M.E.P., K.K., P.S.G., J.E.D., and V.J.J.M. wrote the manuscript with editing help from L.N., L.B., and B.C.

## Additional information

Supplementary information is available in the online version of the paper. Correspondence and requests for materials should be addressed to V.J.J.M.

## ONLINE METHODS

### Plasmids, strains, and growth media

The quadruple auxotrophic *S. cerevisiae* strain BY4741 (*MATa his3Δ1 leu2Δ0 met15Δ0 ura3Δ0*) was employed in this study. The dopamine-producing strain derived from BY4741 (strain 1373) was utilized as the basis for a yeast BIA platform strain. Yeast cultures were grown in YPD medium (10 g L^-1^ Bacto Yeast Extract, 20 g L^-1^ Bacto peptone, 20 g L^-1^ glucose). Transformed cells were selected on YPD agar containing 200 μg mL^-1^ hygromycin B, 400 μg mL^-1^ G418, or a combination of both antibiotics (200 μg mL^-1^ each). Selection using auxotrophic markers was performed in synthetic complete (SC) medium [6.7 g L^-1^ Difco Yeast Nitrogen Base (YNB) without amino acids, 1.62-1.92 g L^-1^ Drop-out Medium Supplements (Millipore-Sigma) minus appropriate amino acids, 20 g L^-1^ glucose). Strains in which prototrophy was restored were selected on YNB medium (6.7 g L^-1^ Difco YNB, 20 g L^-1^ glucose). Plasmids and strains utilized in this work are listed in **Supplementary Tables 3** and **4**, respectively.

### Yeast strain construction

All genetic modifications to yeast were made via CRISPR-Cas9-mediated genomic integration^50,51^ and *in vivo* DNA assembly^52^. Cas9 and gRNA were delivered to yeast using pCas-G418 (ref. ^51^) or a hygromycin-resistance derivative (pCas-Hyg) constructed herein. Linear gRNA cassettes were retargeted by PCR and assembled using *in vivo* gap repair with a linear PCR-generated pCas backbone^50^. Approximately 100 ng of linear pCas was combined with 250 ng of linear gRNA cassette and 500-1,000 ng of total repair DNA in a standard 50 μL lithium acetate transformation. Cells were heat-shocked at 42 °C for 30 minutes, recovered for 16 hours, and plated onto YPD plates containing appropriate antibiotics. Both linear pCas-G418 and pCas-Hyg plasmids were used for transformations involving complex multi-part DNA assemblies. Design of gRNAs^53^ and selection of chromosomal loci for DNA integration^30,54–56^ were carried out according to previous studies. Gene expression cassettes, genomic integration sites, and synthetic DNAs utilized in this work are listed in **Supplementary Tables 5-7**, respectively.

Yeast genes targeted for deletion were replaced with a synthetic DNA landing pad (LP5.T3) possessing a unique Cas9 target site (T3) for subsequent genomic integration of additional copies of the *NdNCSΔN_20_* gene. Seven of the eight copies of *NdNCSΔN_20_* were later deleted in one genome editing event using the *NdNCS* (T_*PGI1*_-LP5) gRNA and a single chromosomal LP5.T3 donor in the *aad3*Δ locus. The remaining *NdNCSΔN_20_* copy was deleted from site FgF20 using an exogenous LP5.T3 donor. *ALD4* was initially deleted using LP5.T3 and reintroduced in later strains along with its cognate promoter and terminator at a different locus (308a).

### Fluorescence microscopy

To visualize GFP-tagged proteins, cells from overnight cultures were back-diluted 50× into fresh SC medium and incubated at 30 °C and 200 rpm for 4-6 hours. Cells were washed with water and mounted unfixed on microscope slides. Images were captured using a Nikon Ti microscope with a 100× PlanAPO lens (NA 1.49). Cells were illuminated using high inclination laminated optical sheet TIRF illumination with 488 nm lasers, and its respective filter cube (Chroma). Images are of single planes. Image processing was done using Fiji (NIH).

### Microtiter plate production assay for dopamine, BIAs, and fusel products

Colonies were picked in triplicate into 0.5 mL of 2× SCS medium (13.4 g L^-1^ Difco Yeast Nitrogen Base (YNB) without amino acids, 2× Drop-out Medium Supplements (Millipore-Sigma) minus appropriate amino acids, 40 g L^-1^ sucrose) within 96-well deep well plates. Following 16-24 h of growth, saturated cultures were back-diluted 50× into 0.5 mL of fresh 2× SCS medium in 96-well deep well plates. Cultures were grown at 30 °C with shaking at 350 rpm. Following 72-96 h of growth, OD_600_ measurements were taken and culture broth was stored at −20 °C for subsequent analysis by LC-MS.

### Cultivation of (*S*)-norcoclaurine and (*S*)-reticuline production strains in a pulsed fed-batch fermentor

Controlled fed-batch fermentations were carried out in 3 L BioBundle fermentors (Applikon). Cultivation temperature was maintained at 30 °C and pH was kept at 4.5 by titration with 4 M NaOH. Dissolved oxygen was maintained at 30% of air saturation by automatically adjusting the stirring rate (aeration rate 1.0 L min^-1^). Off-gas composition (concentration of O_2_ and CO_2_) was analyzed using a Tandem Multiplex gas analyzer (Magellan BioTech). Bioreactor inoculum was generated in two 250 mL shake flasks containing 50 mL of SC-His medium, grown for 24 h at 30 °C. Cells were washed and suspended in 0.9% NaCl, and used to inoculate (OD_600_ = ~0.1) 1 L of batch medium (40 g sucrose, 2.5 g KH_2_PO_4_, 6.0 g (NH_4_)_2_SO_4_, 1.0 g MgSO_4_·7H_2_O, 1.92 g L^-1^ Drop-out Medium Supplements without histidine, 0.076 g L^-1^ l-tryptophan, 0.152 g L^-1^ l-phenylalanine, 5 mL vitamin stock, and 5 mL trace element stock per liter). Vitamin and trace element stock solutions were based on a previous report^57^. The culture was operated in batch mode until sucrose was exhausted (24-30 h), followed by fed-batch phase with automated 10 g L^-1^ sucrose pulses (**Supplementary Fig. 7**). Feeding medium contained 360 g sucrose, 15 g KH_2_PO_4_, 60 g (NH_4_)_2_SO_4_, 6 g MgSO_4_·7H_2_O, 4.16 g l-phenylalanine, 1.55 g l-tryptophan, 15 mL vitamin stock, and 15 mL trace element stock per liter. Samples were collected every 12 hours for a total of five days.

### THIQ synthesis assay from supplemented amino acids

For growth on individual amino acids as the major source of nitrogen, reticuline-producing strain LP501 was transformed with pHUM (ref. ^58^) to complement His and Met auxotrophies. Colonies were first picked in triplicate into 0.5 ml of 2× YNB (without amino acids and ammonium sulfate) containing 40 g L^-1^ sucrose and 1 g L^-1^ urea. Following 24 h of growth, saturated cultures were back-diluted 40× into 0.5 ml of fresh 2× YNB (without amino acids and ammonium sulfate) containing 40 g L^-1^ sucrose within 96-well deep well plates. Cultures of strain LP501 harboring pHUM were also supplemented with 0.076 g L^-1^ each of l-phenylalanine and l-tryptophan due to *pha2*Δ and *trp3*Δ mutations. For THIQ synthesis, cultures were supplemented with individual amino acids to a final concentration of 0.45 g L^-1^ (l-tyrosine), 2 g L^-1^ (l-2-aminoheptanoic acid, l-tryptophan, and l-DOPA), 3 g L^-1^(l-norleucine), or 5 g L^-1^ (l-2-aminobutyrate, l-leucine, l-methionine, l-norvaline, and l-phenylalanine). Ascorbic acid (10 mM) was added to cultures supplemented with l-tryptophan, l-tyrosine, or l-DOPA. Cultures were grown at 30 °C with shaking at 350 rpm. Following 120 h of growth, OD_600_ measurements were taken and culture broth was stored at −20 °C for subsequent analysis by LC-MS.

### LC-MS and HPLC-UV analysis of metabolites

Dopamine, BIA, and other THIQ products from microtiter plate cultures were analyzed using HPLC-FT-MS. Metabolites were extracted from culture broth containing cells and growth medium. For early strains, 25 μL of culture broth was combined with 100 μL of cold 100% acetonitrile (ACN) and 542 μL of 0.123% formic acid was added to give a final concentration of 15% ACN and 0.1% formic acid. For more productive BIA strains and fed-batch cultures, an additional 10-fold dilution of samples was performed. Samples were centrifuged at 4,000 RCF and 10 μL of extracted culture supernatant was separated on a 1290 Infinity II LC system (Agilent Technologies) with a Zorbax Rapid Resolution HT C18 column (30 × 2.1 mm, 1.8 μm; Agilent Technologies). Metabolites were separated using the following gradient: 2% B to 10% B from 0-4 min (0.3 mL min^-1^), 10% B to 85% B from 4-6 min (0.3 mL min^-1^), held at 85% B from 6-7 min (0.3 mL min^-1^), 85%B to 2%B from 7-7.1 min (0.3 mL min^-1^), and held at 2% B from 7.1-9 min (0.45 mL min^-1^). Solvent A was 0.1% formic acid in water and solvent B was 0.1% formic acid in 100% ACN. Following separation, eluent was injected into an LTQ-FT-MS (Thermo Fisher Scientific) using 100 to 400 *m/z* scanning range in positive mode. Resolution, capillary voltage, and source temperature were set to 100,000, 5 kV, and 350 °C, respectively. FT-MS data was processed and manipulated using Xcalibur Qualitative Analysis software (Thermo Fisher Scientific).

Samples producing substituted THIQs from supplemented amino acids were analyzed using an Agilent 6545 quadrupole time-of-flight MS (QTOF-MS; Agilent Technologies) equipped with a Zorbax Eclipse Plus C18 column (50 × 2.1 mm, 1.8 μm; Agilent Technologies) and using the aforementioned gradient conditions. The sample tray and column compartment were set to 4 °C and 30 °C, respectively. The sheath gas flow rate and temperature were adjusted to 10 L min^-1^ and 350 °C, respectively, while drying and nebulizing gases were set to 12 L min^-1^ and 55 psig, respectively. The drying gas temperature was set to 325 °C. QTOF data was processed and manipulated using Agilent MassHunter Qualitative Analysis software.

Dopamine and BIAs from fermentor samples and fusel products from both microtiter plate and fermentor cultures were analyzed and quantified using HPLC-UV according to a modified method^44^. Equal volumes of culture broth and 100% ACN containing 0.1% trifluoroacetic acid (TFA) were combined and samples were centrifuged at 4,000 RCF. Additional dilutions were performed as necessary. Five μL of extracted broth was separated on an Agilent 1200 HPLC system equipped with an Eclipse XDB-C18 column (150 × 4.6 mm, 5 μm, Agilent Technologies). Metabolites were separated using a flow rate of 1 mL min^-1^ and the following gradient: 5% B to 20% B from 0-10 min, 20% B to 50% B from 10-15 min, 50% B to 95% B from 15-15.1 min, and held at 95% B from 15.1-25 min. Solvent A was 0.1% TFA in water and solvent B was 0.1% TFA in 100% methanol. Tyrosol and 4-HPAC were detected at 276 nm; dopamine, (*S*)-norcoclaurine, and (*S*)-reticuline at 280 nm.

