## Supplementary material for "A yeast platform for high-level synthesis of natural and unnatural tetrahydroisoquinoline alkaloids": Pyne et al supplemental information

### Supplementary Results

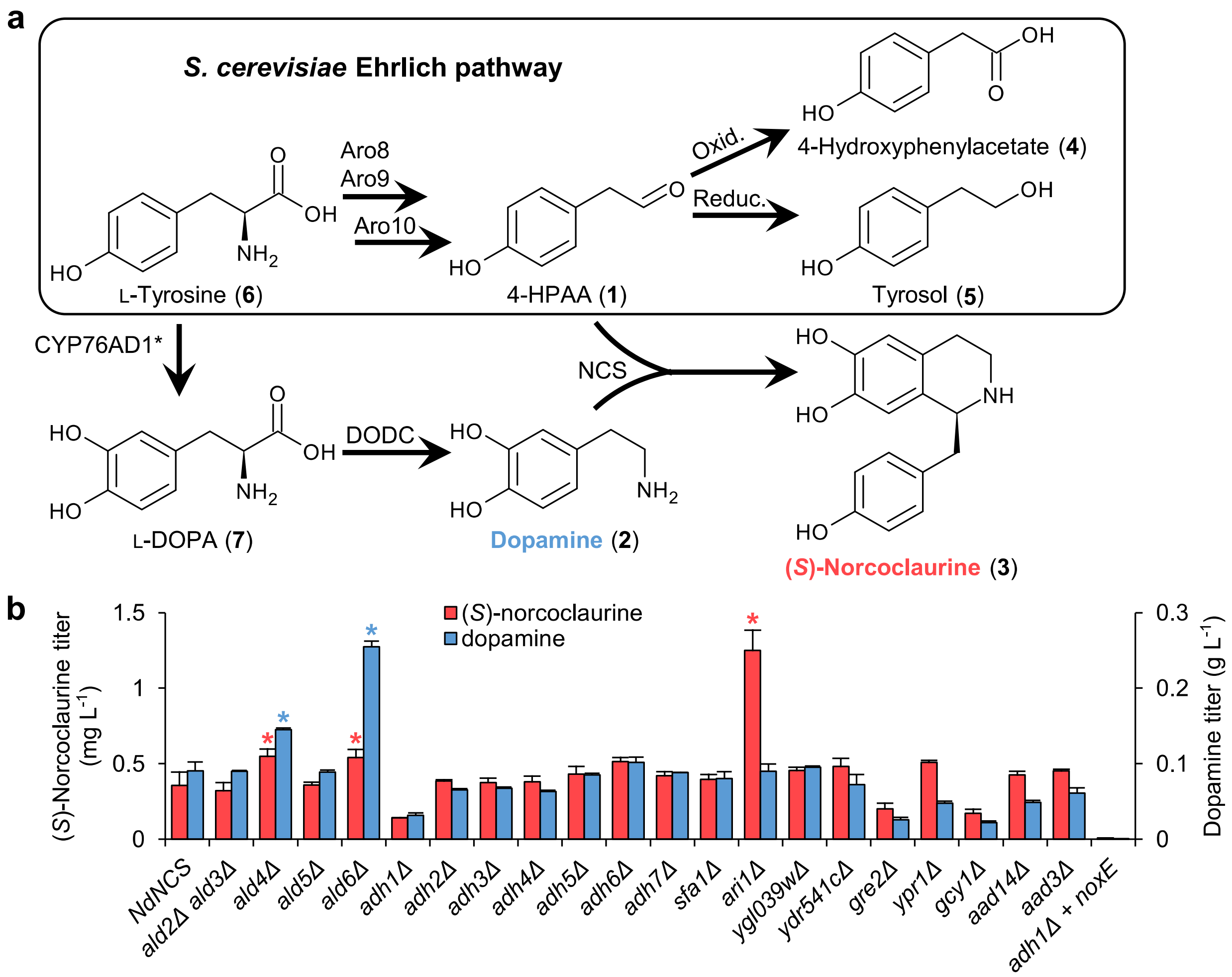


**Supplementary Figure 1. Enhancing 4-HPAA substrate supply through inactivation of host oxidoreductases. (a)** 4-HPAA is produced natively from L-tyrosine via the Ehrlich pathway where it is converted to 4-hydroxyphenylacetate (4-HPAC) or tyrosol. Implementing NCS and a heterologous dopamine biosynthesis pathway diverts 4-HPAA to (*S*)-norcoclaurine formation. (**b**) Dopamine and (*S*)-norcoclaurine titers in culture supernatants of single gene deletion strains as measured by LC-MS. Deletions were performed in a strain harboring a single gene copy of *NdNCS*. *ALD2* and *ALD3* were deleted in conjunction due to proximity in the yeast genome. An NADH oxidase gene (*noxE*) was expressed in the *adh1*Δ mutant to improve growth[1]. Asterisk (*) denotes a significant increase ($P < 0.05$) in titer relative to the parent strain. Error bars represent s.d. of three biological replicates.

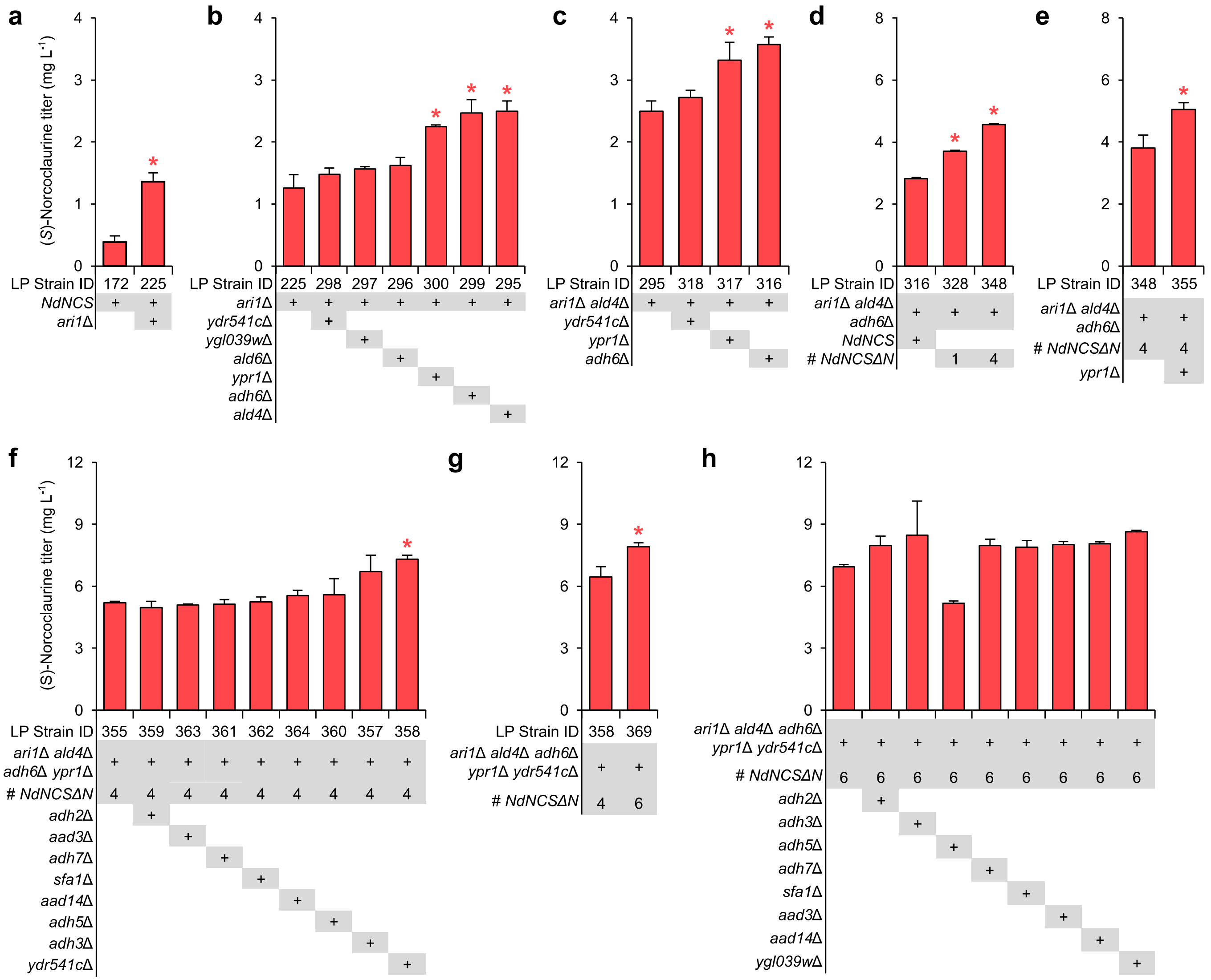


**Supplementary Figure 2. Combinatorial gene deletion analysis to improve (*S*)-norcoclaurine production.** (**a** to **h**) Successive rounds of strain engineering were performed to identify the optimal combination of oxidoreductase gene deletions. NCS activity was also improved through truncation of *NdNCS* (**d**) and increasing copy number of *NdNCSΔN*$_{20}$ (**d** and **g**). Error bars represent s.d. of three biological replicates. Asterisks (*) denote a significant increase ($P < 0.05$) in (*S*)-norcoclaurine production relative to the parent strain.

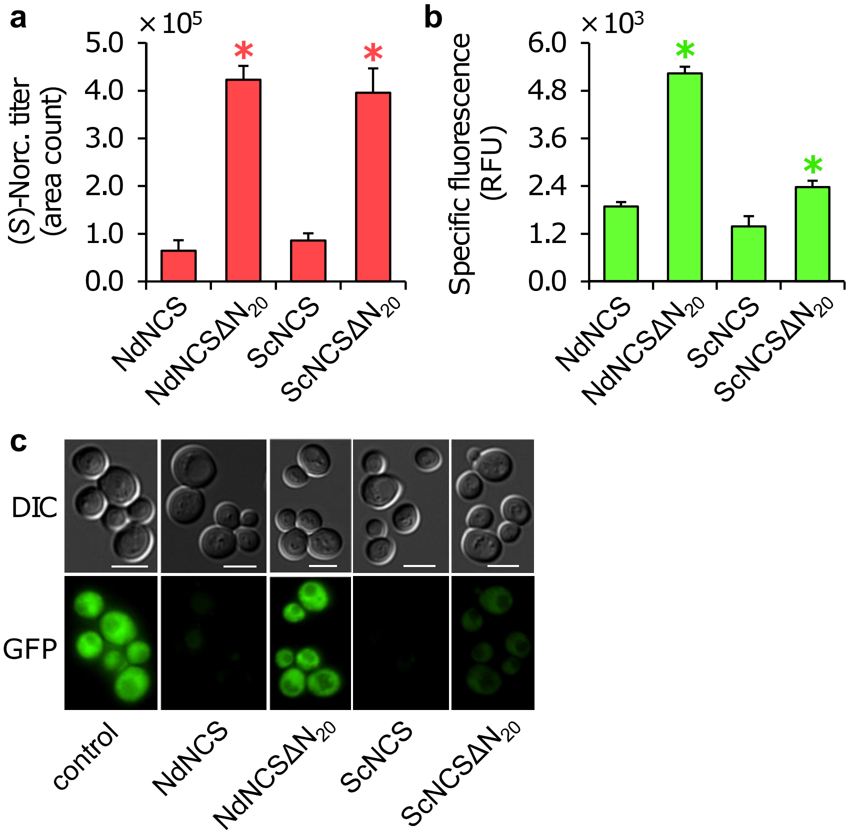


**Supplementary Figure 3. N-Terminal truncation of *Nd*NCS or *Sc*NCS improves production of (*S*)-norcoclaurine.** *Nd*NCS and *Sc*NCS were truncated by removing 20 N-terminal amino acids, yielding *Nd*NCSΔ$N_{20}$ and *Sc*NCSΔ$N_{20}$, respectively. (**a**) (*S*)-Norcoclaurine titer increases following truncation of *Nd*NCS and *Sc*NCS. (**b**) Specific GFP fluorescence (normalized to culture $OD_{600}$) increases following truncation of *Nd*NCS and *Sc*NCS. The C-termini of *Nd*NCS, *Nd*NCSΔ$N_{20}$, *Sc*NCS, and *Sc*NCSΔ$N_{20}$ were fused with GFP. Overnight cultures were back-diluted 50× and grown in 0.5 mL of 2× SC medium for approximately 6 hours. Error bars represent s.d. of three biological replicates. (**c**) N-terminal truncation of *Nd*NCS and *Sc*NCS improves gene expression or enzyme solubility in yeast. Cells of GFP-tagged *Nd*NCS, *Nd*NCSΔ$N_{20}$, *Sc*NCS, and *Sc*NCSΔ$N_{20}$ were visualized using confocal fluorescence microscopy. Control cells harbor GFP without NCS. Scale bars represent 5 µm. Asterisks (*) denote a significant increase ($P < 0.05$) in (*S*)-norcoclaurine production or GFP fluorescence relative to the parent strain.

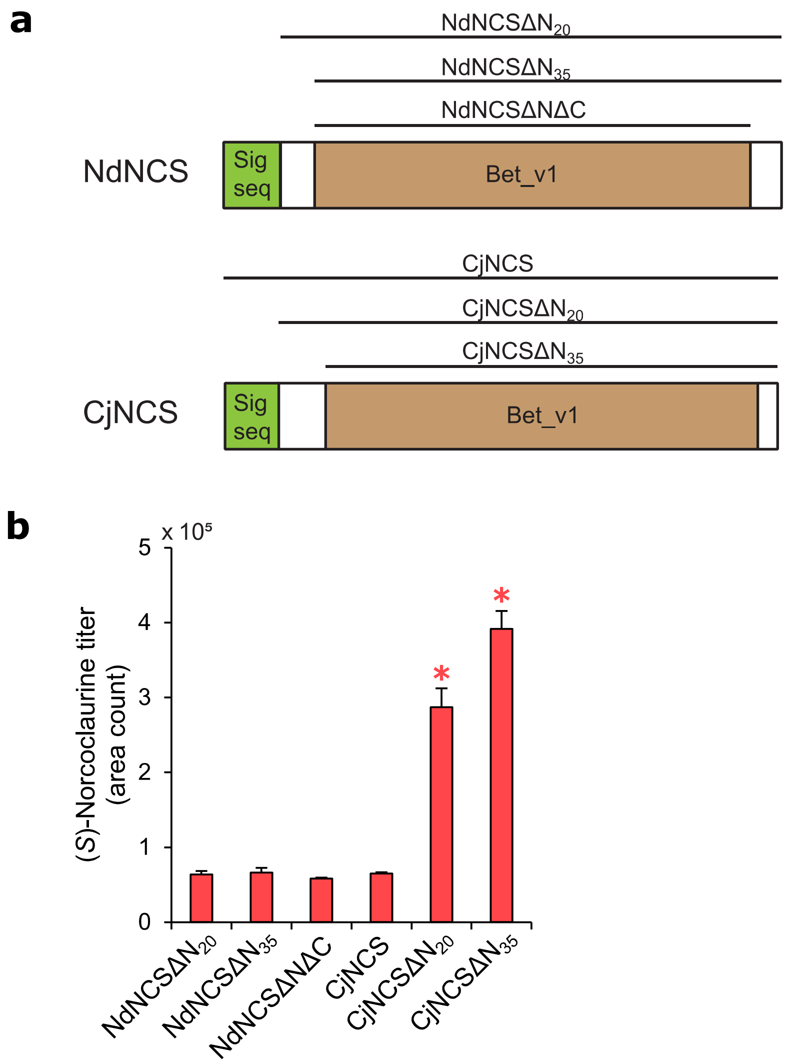


**Supplementary Figure 4. Truncation of *Cj*NCS improves production of (*S*)-norcoclaurine.** *Nd*NCS and *Cj*NCS were truncated by removing various-sized N-terminal and C-terminal regions. (**a**) Structure of N- and C-terminal truncations of *Nd*NCS and *Cj*NCS proteins. Bet_v1 is the putative core NCS catalytic domain required for activity and Sig seq refers to predicted N-terminal signal sequences for targeting *Nd*NCS and *Cj*NCS to subcellular organelles in their respective plant species. (**b**) (*S*)-Norcoclaurine titer increases following truncation of *Cj*NCS to the core Bet_v1 domain. Strains were grown in 0.5 mL of 2× SC medium for 72. Error bars represent s.d. of three biological replicates. Asterisks (*) denote a significant increase ($P < 0.05$) in (*S*)-norcoclaurine production relative to the strain harboring *Nd*NCSΔ$N_{20}$.

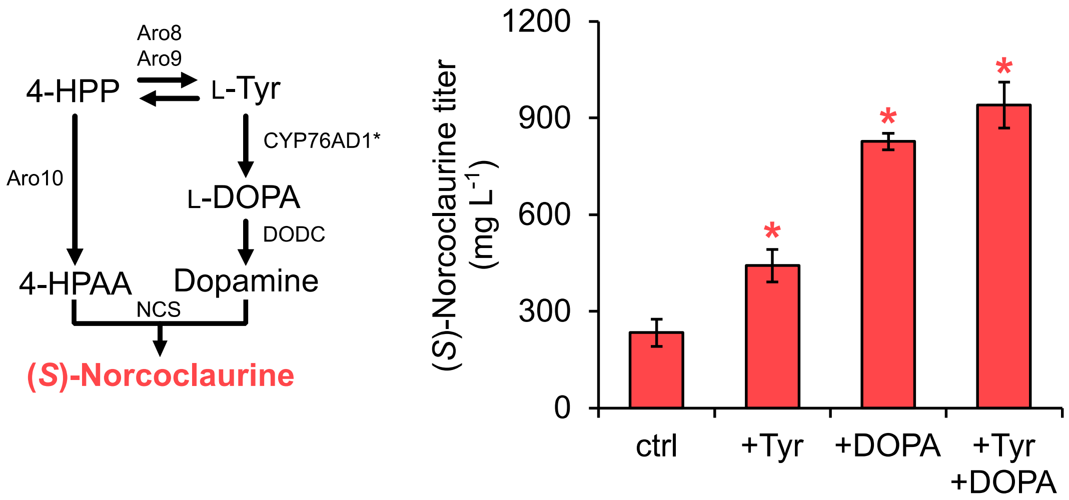


**Supplementary Figure 5. Supplementation of L-DOPA to strain LP412 improves (*S*)-norcoclaurine production.** Exogenously supplied L-DOPA is converted directly to dopamine, whereas L-tyrosine is converted to both 4-HPAA and dopamine. Cultures of strain LP412 were supplemented with 2.5 mM L-tyrosine, 5 mM L-DOPA, or a combination of both amino acids and grown in 0.5 mL of 2× SC medium for 72 hours. Ten mM sodium ascorbate was added to all cultures to limit oxidation of aromatic amino acids. Error bars represent s.d. of four biological replicates. Asterisks (*) denote a significant increase ($P < 0.05$) in (*S*)-norcoclaurine production relative to the control culture.

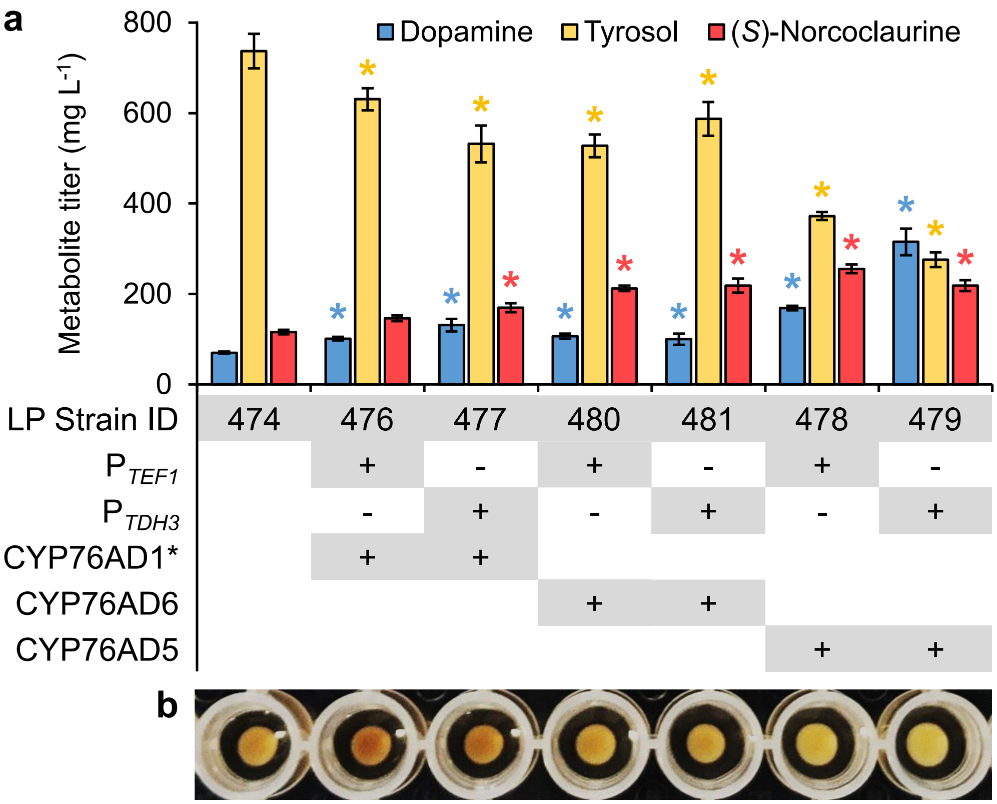


**Supplementary Figure 6. Expression of *CYP76AD5* or *CYP76AD6* enhances (*S*)-norcoclaurine biosynthesis.** Tyrosine hydroxylase variants (*CYP76AD1**, *CYP76AD6*, or *CYP76AD5*) were integrated into strain LP474 and expressed from one of two promoters ($P_{TEF1}$ or $P_{TDH3}$). Strain LP474 and its derivatives contain an existing copy of $P_{TDH3}$-*CYP76AD1** (*CYP76AD1*$^{W13L\ F309L}$). (**a**) Implementation of *CYP76AD5*, *CYP76AD6*, or an additional copy of *CYP76AD1** yields a range of tyrosol, dopamine, and (*S*)-norcoclaurine titers. CYP76AD5 is a more active enzyme than CYP76AD1* and CYP76AD6 (ref. [2]). $P_{TDH3}$ is a stronger promoter than $P_{TEF1}$ (ref. [3,4]). Expression of *CYP76AD5* from $P_{TEF1}$ yielded the highest (*S*)-norcoclaurine titer of all strains assayed, while its expression from $P_{TDH3}$ yielded the highest dopamine titer and the lowest concentration of tyrosol. Error bars represent s.d. of three biological replicates. Asterisks (*) denote a significant increase or decrease ($P < 0.05$) in metabolite production relative to strain LP474. (**b**) Pigmentation of cells expressing *CYP76AD1**, *CYP76AD6*, or *CYP76AD5*. CYP76AD1 and its engineered variant (CYP76AD1*) possess DOPA oxidase side activity not observed in CYP76AD5 and CYP76AD6 (ref. [2]), which results in the accumulation of melanin, a brown pigment[5]. Strains for pigmentation and metabolite production assays were grown in 0.5 mL of 2× SC medium for 96 hours.

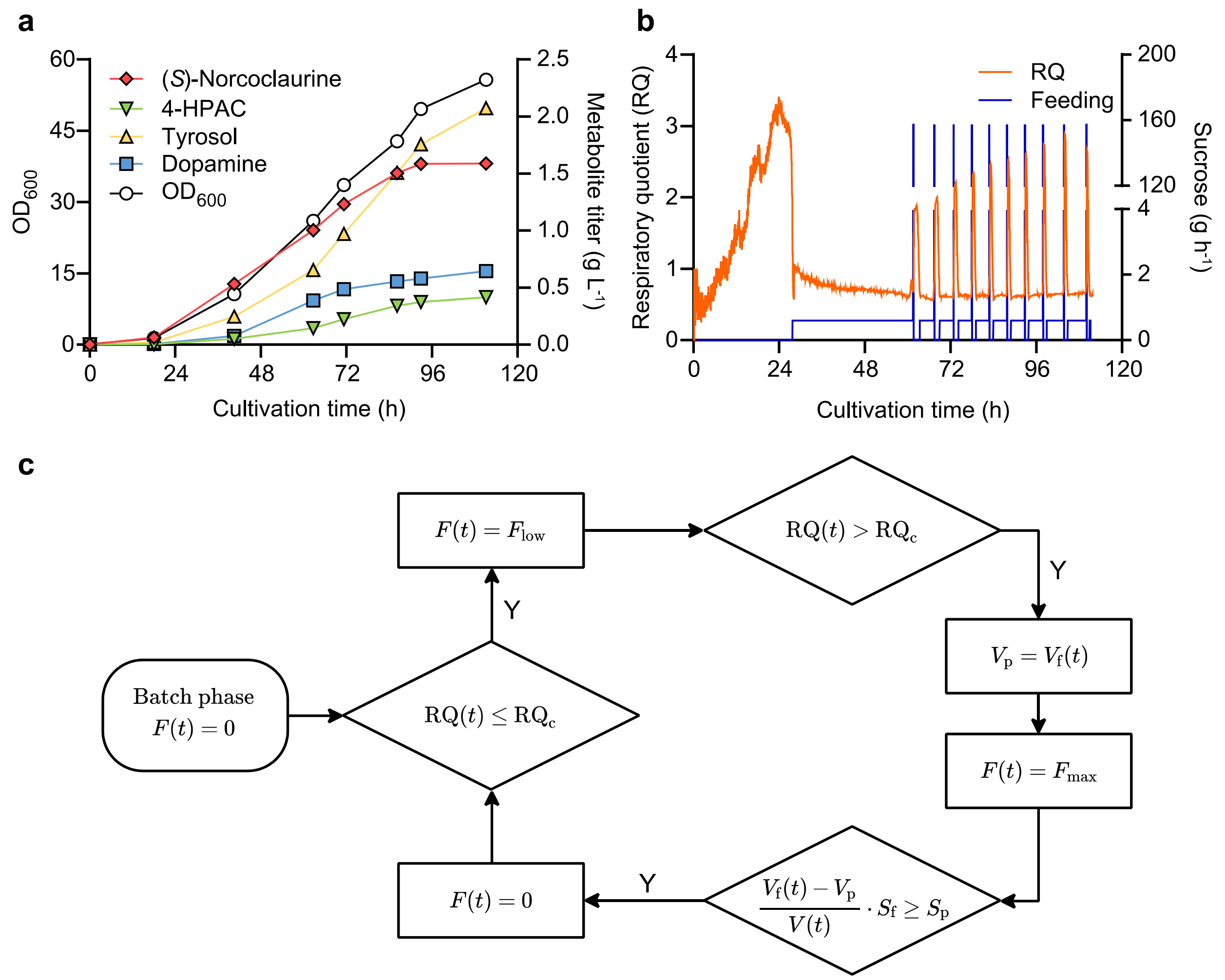


**Supplementary Figure 7. Cultivation of an (*S*)-norcoclaurine-producing strain (LP478) in a sucrose-pulsed fed-batch fermentor.** (**a**) Growth of biomass ($OD_{600}$) and accumulation of BIA metabolites (measured using HPLC-UV) in the culture medium during cultivation. (**b**) Cells were grown in batch phase until exhaustion of sucrose (40 g $L^{-1}$), indicated by a rapid drop of respiratory quotient (RQ) value, triggering constant feeding of fed-batch medium (corresponding to 0.60 g $h^{-1}$ sucrose). Constant feeding continued until the exhaustion of ethanol (indicated by an increase in RQ value) produced in the batch phase. Subsequently, a pulse of fed-batch medium (corresponding to 10 g $L^{-1}$ sucrose) was rapidly fed into the reactor, after which the pump was stopped. Feeding (0.60 g $h^{-1}$ sucrose) resumed after exhaustion of sucrose, and until consumption of ethanol, after which another pulse (10 g $L^{-1}$ sucrose) was fed, continuing the cycle. (**c**) Logic chart of pulse feeding algorithm. $F(t)$, substrate (sucrose) feeding rate at time $t$; $RQ(t)$, on-line value of RQ at time $t$; $RQ_c$, trigger value of RQ; $F_{low}$, substrate feeding rate setpoint for constant feeding; $F_{max}$, substrate feeding rate setpoint for pulse feeding; $V_f(t)$, volume of feeding medium fed into the reactor at time $t$; $V_p$, feeding volume storage value; $V(t)$, current culture volume; $S_f$, concentration of sucrose in the feeding medium; $S_p$, target concentration of sucrose in the bioreactor after a pulse; $t$, process time.

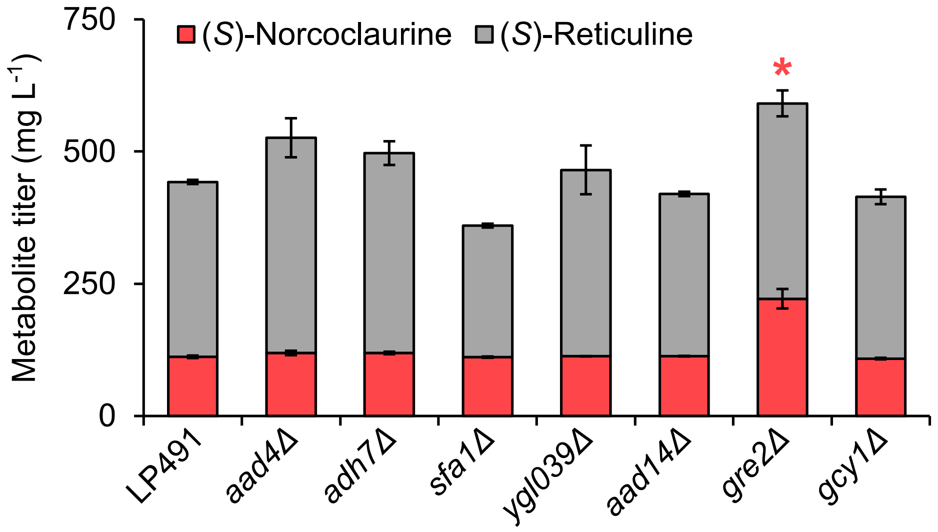


**Supplementary Figure 8. Deletion of *GRE2* in strain LP491 increases THIQ production in microtiter plate cultures.** (*S*)-Norcoclaurine and (*S*)-reticuline titers in culture supernatants of strain LP491 containing deletions in oxidoreductase genes. Strain LP491 is an (*S*)-reticuline-producing strain containing deletions in five oxidoreductase genes (*ari1*Δ *adh6*Δ *ypr1*Δ *ydr541c*Δ *aad3*Δ). Deletion of *GRE2* facilitates a significant increase in (*S*)-norcoclaurine rather than (*S*)-reticuline production due to a presumed bottleneck in an (*S*)-reticuline pathway enzyme in microtiter plate cultures. Asterisk (*) denotes a significant increase ($P < 0.05$) in (*S*)-norcoclaurine titer relative to strain LP491. Error bars represent s.d. of three biological replicates.

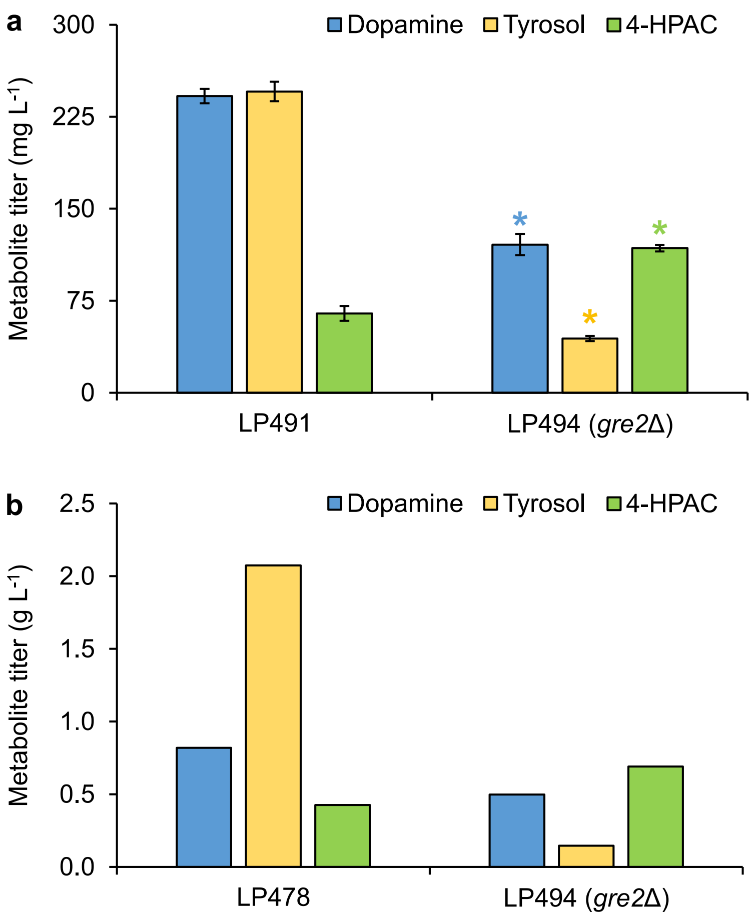


**Supplementary Figure 9. Deletion of *GRE2* diminishes tyrosol synthesis in microtiter plate and pulsed fed-batch fermentor cultures.** (**a**) Dopamine and fusel product synthesis in microtiter plate cultures. Strains LP491 and LP494 harbor deletions in five oxidoreductase genes (*ari1*Δ *adh6*Δ *ypr1*Δ *ydr541c*Δ *aad3*Δ), while LP494 contains an additional deletion in the *GRE2* gene, resulting in reduced levels of dopamine and tyrosol, and an increase in 4-HPAC concentration. Error bars represent s.d. of three biological replicates. Asterisks (*) denote a significant increase or decrease ($P < 0.05$) in metabolite production relative to strain LP491. (**b**) Dopamine and fusel product synthesis in pulsed fed-batch fermentor cultures. Strain LP478 harbors deletions in five oxidoreductases (*ari1*Δ *adh6*Δ *ypr1*Δ *ydr541c*Δ *aad3*Δ), while LP494 contains an additional deletion in *GRE2*. Data is shown from the samples possessing the highest concentration of tyrosol from single fermentor experiments.

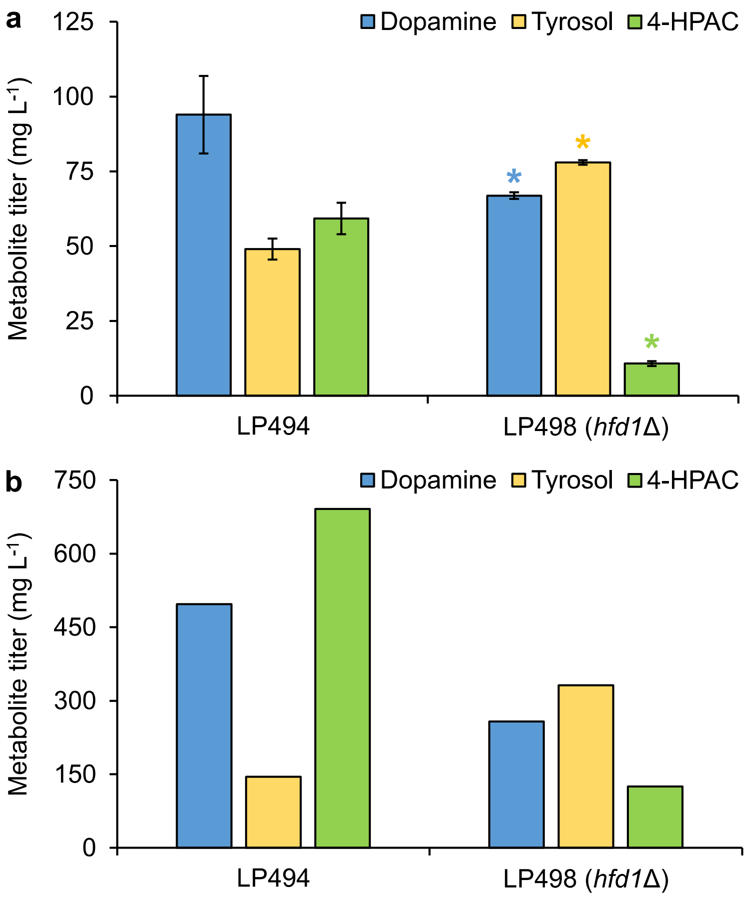


**Supplementary Figure 10. Deletion of *HFD1* diminishes 4-HPAC synthesis in microtiter plate and pulsed fed-batch fermentor cultures.** (**a**) Dopamine and fusel product synthesis in microtiter plate cultures. Strains LP494 and LP498 harbor deletions in six oxidoreductase genes (*ari1*Δ *adh6*Δ *ypr1*Δ *ydr541c*Δ *aad3*Δ *gre2*Δ), while LP498 contains an additional deletion in the *HFD1* aldehyde dehydrogenase gene. Deletion of *HFD1* results in reduced levels of dopamine and 4-HPAC. Error bars represent s.d. of three biological replicates. Asterisks (*) denote a significant increase or decrease ($P < 0.05$) in metabolite production relative to strain LP494. (**b**) Dopamine and fusel product synthesis in pulsed fed-batch fermentor cultures. Data is shown from the samples possessing the peak concentration of 4-HPAC from single fermentor experiments.

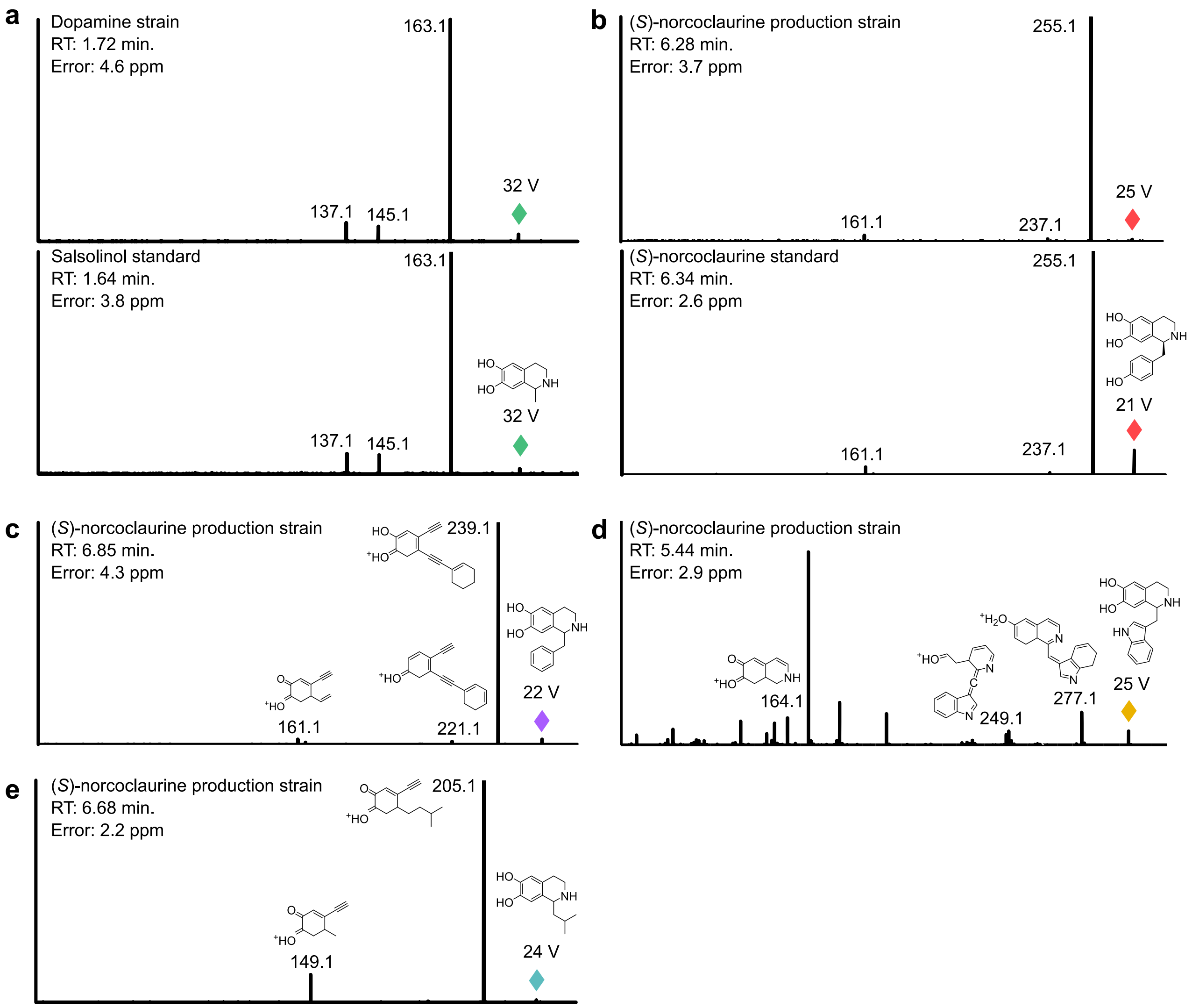


**Supplementary Figure 11. Fragmentation spectra of substituted tetrahydroisoquinoline structures synthesized *de novo*.** (**a**) Salsolinol (**13**). (**b**) (*S*)-Norcoclaurine (**3**). (**c**) Product **16**. (**d**) Product **19**. (**e**) Product **22**. Parent ions are depicted in color and collision energies (V) and mass errors (ppm) are shown. Fragmentation spectra of salsolinol (**13**) and (*S*)-norcoclaurine (**3**) were compared to spectra of authentic standards. Other structures were modelled using the CFM-ID tool[6]. Fragment structures and exact masses are shown for peaks that were matched using CFM-ID. Several observed peaks of **19** could not be matched with predicted fragments. It has been reported that indole-containing molecules undergo complex rearrangements upon fragmentation[7]. Stereochemistry of unnatural substituted tetrahydroisoquinolines is omitted. Salsolinol (**13**), (*S*)-norcoclaurine (**3**), **16**, and **22** were analyzed using FT-MS/MS. Product **19** was analyzed using QTOF-MS/MS. Repeating MS/MS fragmentations of all structures routinely yielded similar results.

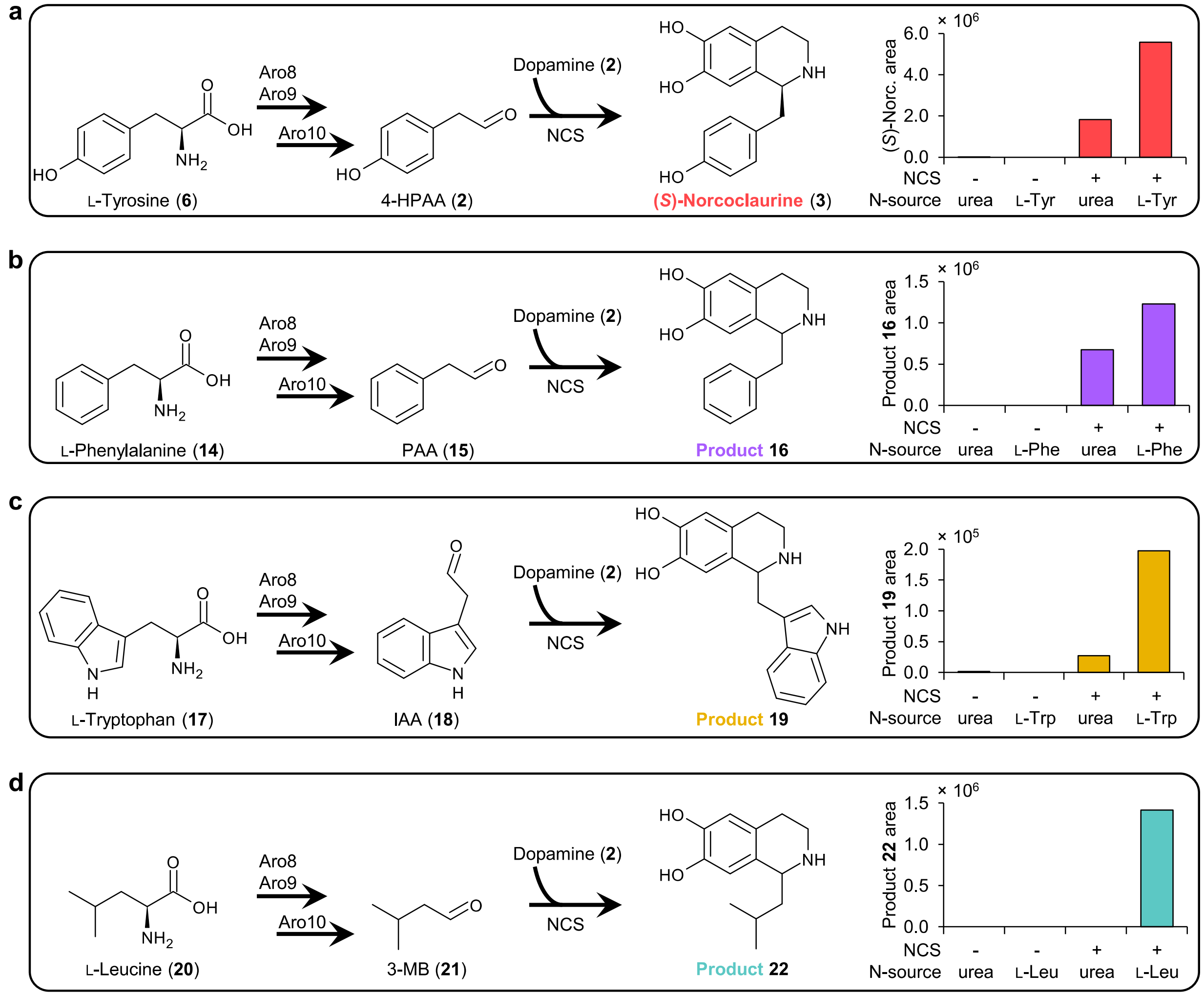


**Supplementary Figure 12. Unnatural substituted tetrahydroisoquinolines derive from amino acids.** A dopamine-producing strain harboring *Cj*NCSΔN$_{35}$ (strain LP385) was cultivated on urea or Ehrlich pathway amino acids as a sole source of nitrogen. (**a**) Synthesis of (*S*)-norcoclaurine (**3**) increases upon growth on L-tyrosine (**6**). (**b**) Synthesis of **16** increases upon growth on L-phenylalanine (**14**). (**c**) Synthesis of **19** increases upon growth on L-tryptophan (**17**). (**d**) Growth on L-leucine (**20**) is essential to observe formation of **22** using strain LP385. Stereochemistry of non-canonical substituted tetrahydroisoquinolines is omitted. Repeating supplementation experiments routinely yielded similar results. Abbreviations: 4-HPAA, 4-hydroxyphenylacetaldehyde; IAA, indole acetaldehyde; 3-MB, 3-methylbutanal; PAA, phenylacetaldehyde; spont., spontaneous.

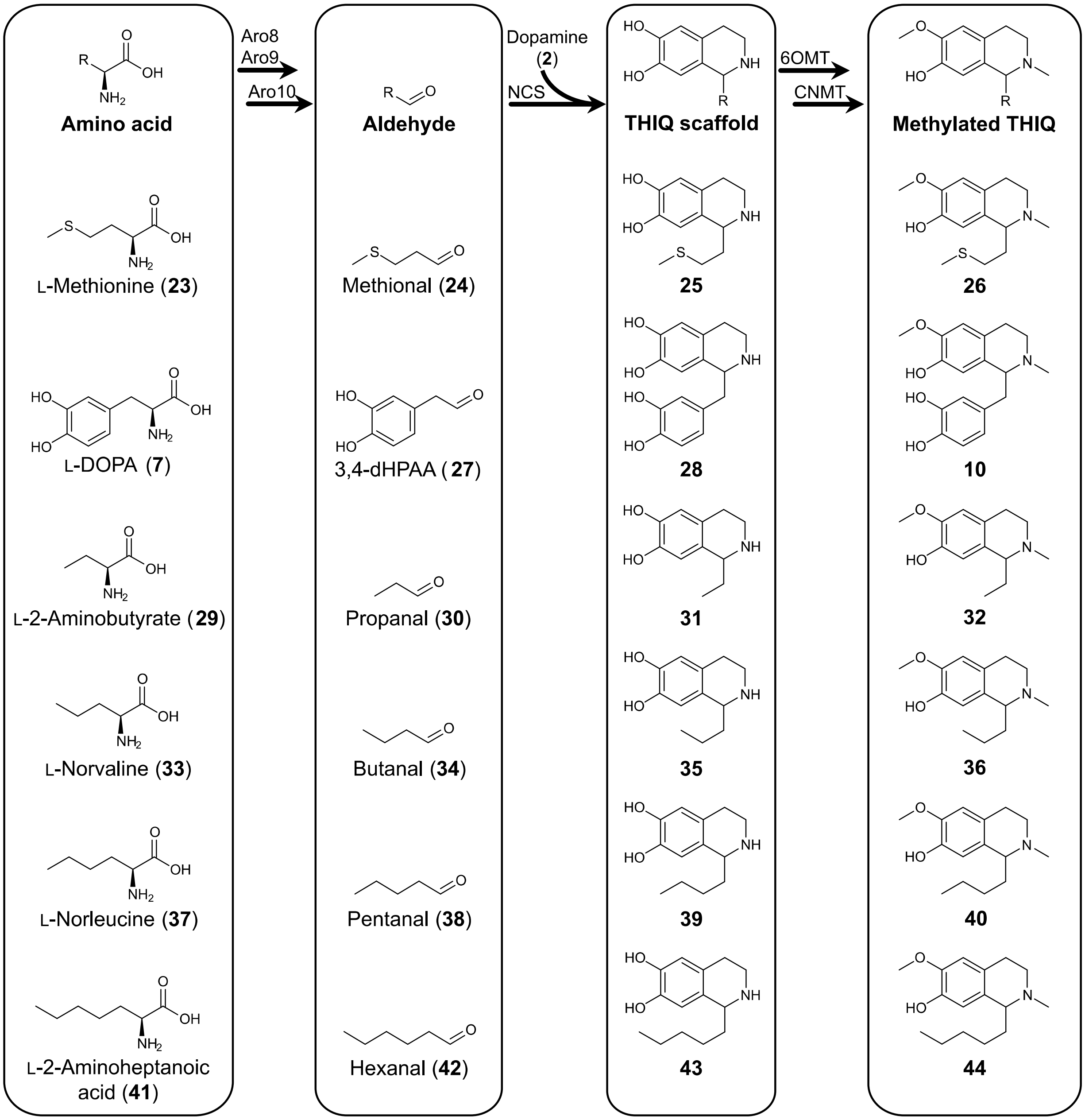


**Supplementary Figure 13. Structures of substituted tetrahydroisoquinolines (THIQs) synthesized from supplemented amino acids.** Amino acids are catabolized to the respective aldehyde species via the yeast Ehrlich pathway (Aro8/Aro9 + Aro10). In the presence of dopamine (**2**) and *Cj*NCSΔ$N_{35}$ (strain LP501), aldehydes are diverted to THIQ synthesis. Strain LP501 contains *Ps*6OMT and *Ps*CNMT methyltransferases for decorating THIQ scaffolds. Stereochemistry of substituted THIQs is omitted.

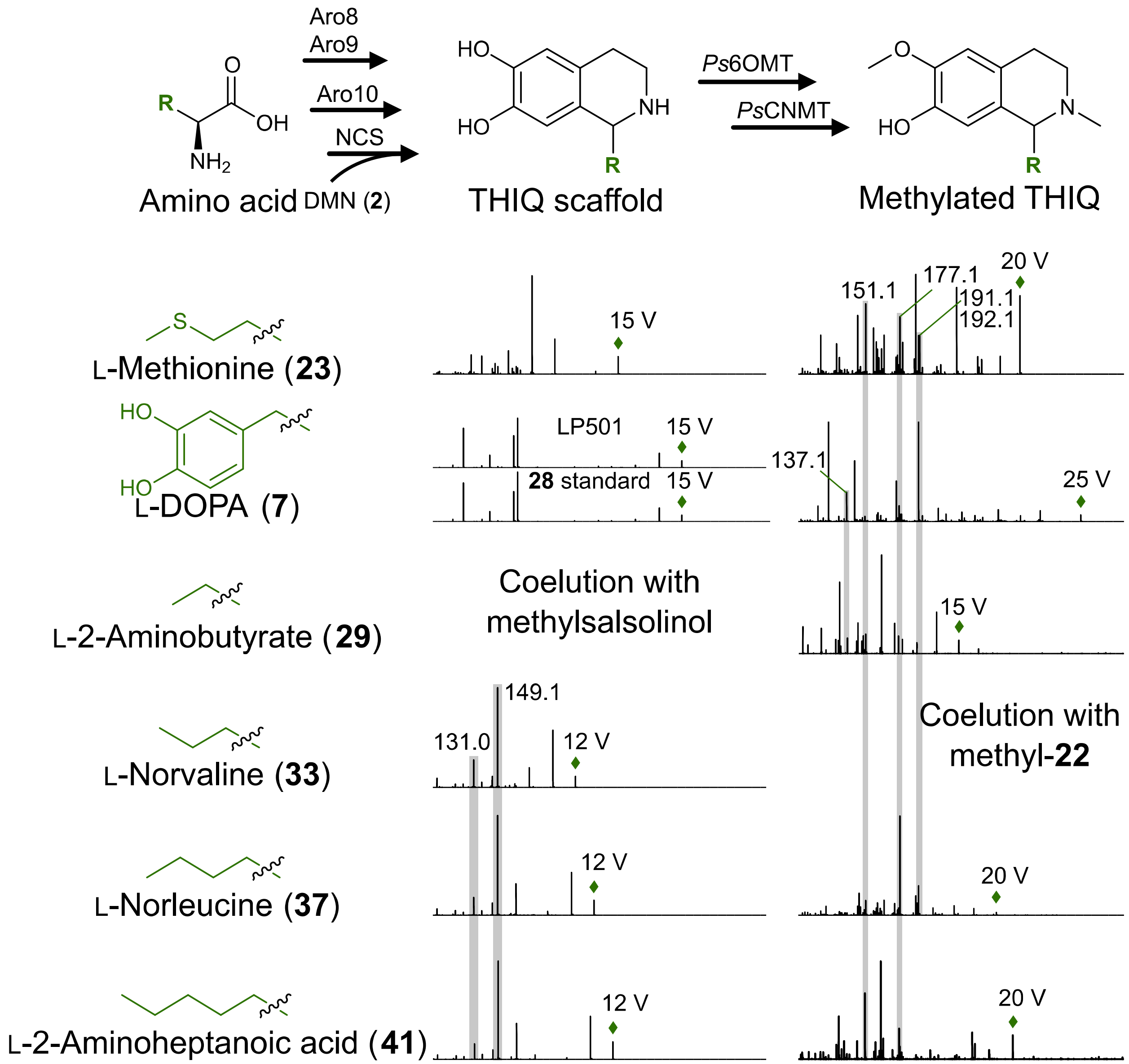


**Supplementary Figure 14. Fragmentation spectra of substituted tetrahydroisoquinoline structures synthesized from supplemented amino acids.** Parent ions are depicted with a green diamond. Fragmentation spectra of norlaudanosoline (**28**) was compared to that of authentic standard. Other structures were modelled using the CFM-ID tool[6]. Masses are shown for key peaks corresponding to THIQ fragments. Unmethylated alkyl-substituted tetrahydroisoquinolines yield characteristic fragments of *m/z* 131.0 and 149.1. Fragment ions of *m/z* 137.1, 151.1, 177.1, and 191.1 possess the 6OMT modification and *m/z* 192.1 possesses both 6OMT and CNMT modifications. Repeating MS/MS fragmentations routinely yielded similar results.

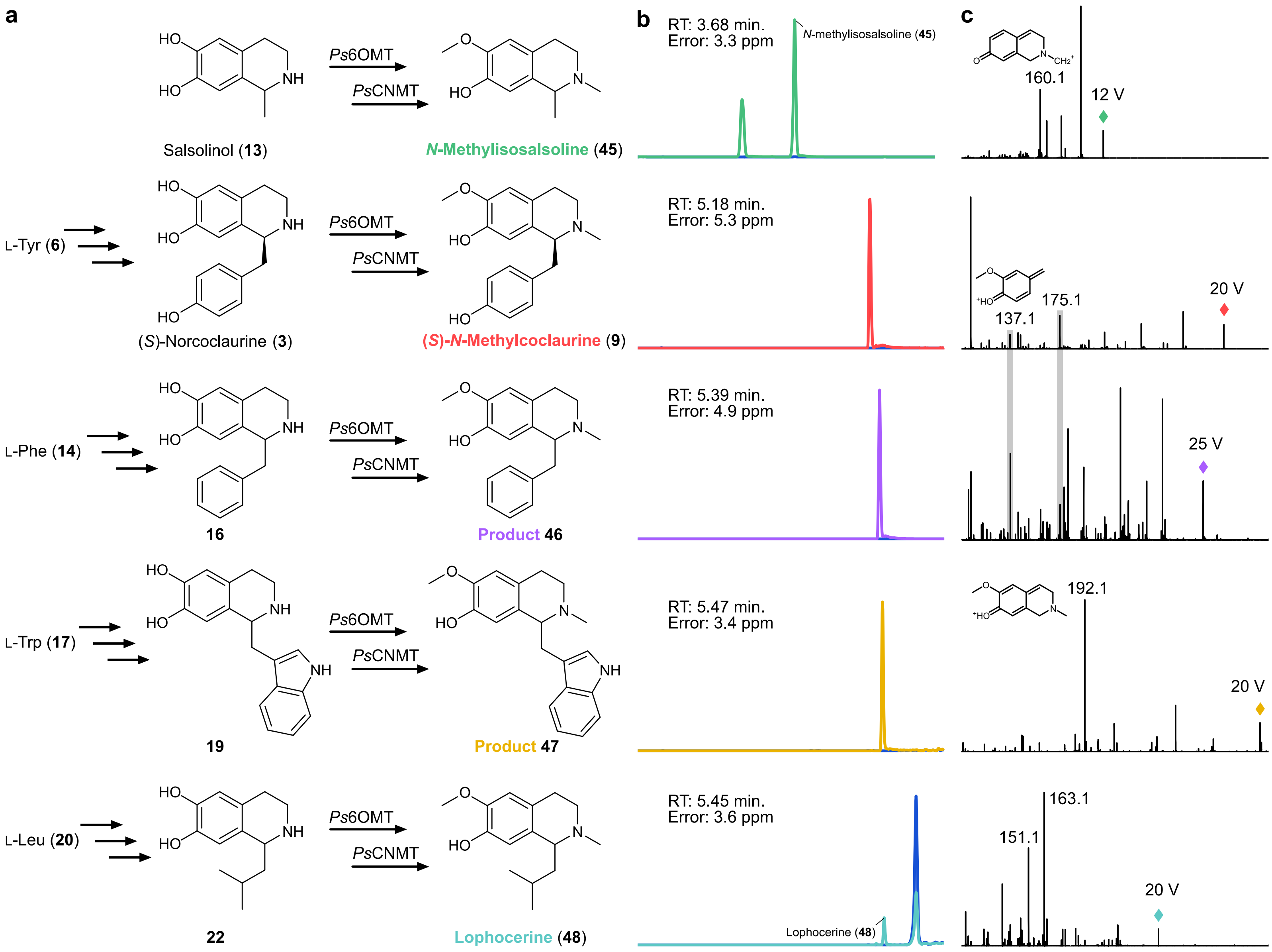

a
b
c
Salsolinol (13)
N-Methylisosalsoline (45)
L-Tyr (6)
(S)-Norcoclaurine (3)
(S)-N-Methylcoclaurine (9)
L-Phe (14)
16
Product 46
L-Trp (17)
19
Product 47
L-Leu (20)
22
Lophocerine (48)
Ps6OMT
PsCNMT
RT: 3.68 min.
Error: 3.3 ppm
N-methylisosalsoline (45)
RT: 5.18 min.
Error: 5.3 ppm
RT: 5.39 min.
Error: 4.9 ppm
RT: 5.47 min.
Error: 3.4 ppm
RT: 5.45 min.
Error: 3.6 ppm
Lophocerine (48)
160.1
12 V
137.1
175.1
20 V
25 V
192.1
20 V
163.1
151.1
20 V

**Supplementary Figure 15.** ***De novo*** **synthesis of methylated substituted tetrahydroisoquinolines (THIQs) by an (*****S*****)-reticuline production host (strain LP501).** (**a**) Decoration of *de novo* THIQ scaffolds by (*S*)-reticuline-pathway methyltransferases. Salsolinol (**13**), (*S*)-norcoclaurine (**3**), **16**, **19**, and **22** are synthesized from endogenous acetaldehyde (**12**), L-tyrosine (**6**), L-phenylalanine (**14**), L-tryptophan (**17**), and L-leucine (**20**), respectively. Strain LP501 produces *Ps*6OMT and *Ps*CNMT methyltransferases for decorating THIQ scaffolds, yielding *N*-methylisosalsoline (**45**), (*S*)-*N*-methylcoclaurine (**9**), **46**, **47**, and lophocerine (**48**). Stereochemistry of non-canonical substituted THIQs is omitted. (**b**) Ion-extracted LC-QTOF-MS chromatograms of strain LP501 grown on urea. Methylated substituted THIQs shifted in retention time (RT) relative to the canonical methylated product from L-tyrosine [(*S*)-*N*-methylcoclaurine (**9**)]. Growth of a dopamine-producing strain that lacks NCS, 6OMT, and CNMT enzymes (strain 1373) under the same conditions (blue) failed to generate peaks corresponding to substituted THIQs. All *m/z* values were calculated based on the expected theoretical structure of the respective compounds of interest and mass error (ppm) is shown. (**c**) QTOF-MS/MS fragmentation spectra of methylated substituted THIQ structures synthesized *de novo*. Parent ions are depicted using colored diamonds and collision energies are shown. Fragmentation spectra were modelled using the CFM-ID tool[6]. Structures and/or masses are shown for key peaks corresponding to methylated THIQ fragments. Fragments of *m/z* 137.1, 151.1, and 163.1 possess the 6OMT modification, *m/z* 160.1 contains the CNMT modification, and *m/z* 192.1 possesses both 6OMT and CNMT methyl groups. Fragments of *m/z* 137.1, 175.1, and 192.1 derive from fragmentation of (*S*)-reticuline[5]. Repeating MS/MS fragmentations of all structures routinely yielded similar results.

**Table S1 – Putative substituted tetrahydroisoquinoline products queried using FT-MS**

| Carbonyl[a] | Carbonyl formula | THIQ formula | Expected THIQ mass (*m/z* + H$^+$) | Observed THIQ mass (*m/z* + H$^+$) | Mass error (ppm) |
|---|---|---|---|---|---|
| *Aliphatic aldehydes* | | | | | |
| Acetaldehyde | $C_2H_4O$ | $C_{10}H_{13}NO_2$ (Salsolinol, **13**) | 180.1024 | 180.1018 | 3.3 |
| Butanal | $C_4H_8O$ | $C_{12}H_{17}NO_2$ | 208.1338 | - | |
| Methional | $C_4H_8OS$ | $C_{12}H_{17}NO_2S$ | 240.1058 | - | |
| 3-Methylbutanal | $C_5H_{10}O$ | $C_{13}H_{19}NO_2$ (**22**) | 222.1494 | 222.1487 | 3.2 |
| 2-Methylbutanal[b] | $C_5H_{10}O$ | $C_{13}H_{19}NO_2$ | 222.1494 | - | - |
| 2-Methylpropanal | $C_4H_8O$ | $C_{12}H_{17}NO_2$ | 208.1338 | - | |
| 2-Octenal | $C_8H_{14}O$ | $C_{16}H_{23}NO_2$ | 262.1807 | - | |
| 2-Decenal | $C_{10}H_{18}O$ | $C_{18}H_{27}NO_2$ | 290.2120 | - | |
| *Aromatic aldehydes* | | | | | |
| Benzaldehyde | $C_7H_6O$ | $C_{15}H_{15}NO_2$ | 242.1181 | - | |
| 2-Hydroxy benzaldehyde | $C_7H_6O_2$ | $C_{15}H_{15}NO_3$ | 258.1130 | - | |
| 2-Phenylacetaldehyde | $C_8H_8O$ | $C_{16}H_{17}NO_2$ (**16**) | 256.1338 | 256.1333 | 2.0 |
| 3,4-Dihydroxyphenylacetaldehyde | $C_8H_8O_3$ | $C_{16}H_{17}NO_4$ | 288.1236 | - | |
| Phenylpyruvate | $C_9H_8O_3$ | $C_{17}H_{17}NO_4$ | 300.1236 | - | |
| Hydroxyphenylpyruvate | $C_9H_8O_4$ | $C_{17}H_{17}NO_5$ | 316.1185 | - | |
| 3,5-Dimethyl-benzaldehyde | $C_9H_{10}O$ | $C_{17}H_{19}NO_2$ | 270.1494 | - | |
| Syringaldehyde | $C_9H_{10}O_4$ | $C_{17}H_{19}NO_5$ | 318.1342 | - | |
| Indole-3-acetaldehyde | $C_{10}H_9NO$ | $C_{18}H_{18}N_2O_2$ (**19**) | 295.1446 | 295.1442 | 1.4 |
| 2-Phenyl-2-butenal | $C_{10}H_{10}O$ | $C_{18}H_{20}NO_2$ | 283.1572 | - | |
| *Aliphatic ketones* | | | | | |
| 2-Butanone | $C_4H_8O$ | $C_{12}H_{17}NO_2$ | 208.1338 | - | |
| 2,3-Butanedione | $C_4H_6O_2$ | $C_{12}H_{15}NO_3$ | 222.1130 | - | |

| | | | | |
|---|---|---|---|---|
| 2,3-Pentanedione | $C_5H_8O_2$ | $C_{13}H_{17}NO_3$ | 236.1287 | - |
| 2-Pentanone | $C_5H_{10}O$ | $C_{13}H_{19}NO_2$ | 222.1494 | - |
| 3-Hydroxy-2-pentanone | $C_5H_{10}O_2$ | $C_{13}H_{19}NO_3$ | 238.1443 | - |
| 4-Methyl-3-penten-2-one | $C_6H_{10}O$ | $C_{14}H_{19}NO_2$ | 234.1494 | - |
| 2,5-Hexanedione | $C_6H_{10}O_2$ | $C_{14}H_{19}NO_3$ | 250.1443 | - |
| 4-Methyl-2-pentanone | $C_6H_{12}O$ | $C_{14}H_{21}NO_2$ | 236.1650 | - |
| 2-Hexanone | $C_6H_{12}O$ | $C_{14}H_{21}NO_2$ | 236.1650 | - |
| 4-Hydroxy-4-methyl-2-pentanone | $C_6H_{12}O_2$ | $C_{14}H_{21}NO_3$ | 252.1600 | - |
| 2,3-Heptanedione | $C_7H_{12}O_2$ | $C_{15}H_{21}NO_3$ | 264.1600 | - |
| 4-Methyl-2-hexanone | $C_7H_{14}O$ | $C_{15}H_{23}NO_2$ | 250.1807 | - |
| 5-Methyl-2-hexanone | $C_7H_{14}O$ | $C_{15}H_{23}NO_2$ | 250.1807 | - |
| 2-Heptanone | $C_7H_{14}O$ | $C_{15}H_{23}NO_2$ | 250.1807 | - |
| 4-Methyl-3-hepten-2-one (isomer) | $C_8H_{14}O$ | $C_{16}H_{23}NO_2$ | 262.1807 | - |
| 4-Methyl-3-hepten-2-one (isomer) | $C_8H_{14}O$ | $C_{16}H_{23}NO_2$ | 262.1807 | - |
| 2,3-Octanedione | $C_8H_{14}O_2$ | $C_{16}H_{23}NO_3$ | 278.1756 | - |
| 3,4-Dimethyl-2-hexanone | $C_8H_{16}O$ | $C_{16}H_{25}NO_2$ | 264.1964 | - |
| 6-Methyl-2-heptanone | $C_8H_{16}O$ | $C_{16}H_{25}NO_2$ | 264.1964 | - |
| 4-Methyl-2-heptanone | $C_8H_{16}O$ | $C_{16}H_{25}NO_2$ | 264.1964 | - |
| 2,5-Dimethyl-3-hexanone | $C_8H_{16}O$ | $C_{16}H_{25}NO_2$ | 264.1964 | - |
| 4-Octanone | $C_8H_{16}O$ | $C_{16}H_{25}NO_2$ | 264.1964 | - |
| 2,6-Dimethyl-2,5-heptadien-4-one | $C_9H_{14}O$ | $C_{17}H_{23}NO_2$ | 274.1807 | - |
| 2-Methyl-2-octen-4-one | $C_9H_{16}O$ | $C_{17}H_{25}NO_2$ | 276.1964 | - |
| 5-Nonen-2-one | $C_9H_{16}O$ | $C_{17}H_{25}NO_2$ | 276.1964 | - |
| 2,4-Dimethyl-3-heptanone | $C_9H_{18}O$ | $C_{17}H_{27}NO_2$ | 278.2120 | - |
| C9 Ketone (*m/z* 43, 57, 86) | $C_9H_{18}O$ | $C_{17}H_{27}NO_2$ | 278.2120 | - |
| C9 Ketone (*m/z* 57, 72, 99) | $C_9H_{18}O$ | $C_{17}H_{27}NO_2$ | 278.2120 | - |
| C9 Ketone (*m/z* 43, 71) | $C_9H_{18}O$ | $C_{17}H_{27}NO_2$ | 278.2120 | - |
| C9 Ketone (*m/z* 43, 58, 85) | $C_9H_{18}O$ | $C_{17}H_{27}NO_2$ | 278.2120 | - |
| 5-Nonanone | $C_9H_{18}O$ | $C_{17}H_{27}NO_2$ | 278.2120 | - |
| 2-Nonanone | $C_9H_{18}O$ | $C_{17}H_{27}NO_2$ | 278.2120 | - |
| 3,3,6-Trimethylhepta-1,5-dien-4-one | $C_{10}H_{16}O$ | $C_{18}H_{25}NO_2$ | 288.1964 | - |
| 2-Methyl-2-nonen-4-one | $C_{10}H_{18}O$ | $C_{18}H_{27}NO_2$ | 290.2120 | - |
| 3,5-Dimethyl-2-octanone | $C_{10}H_{20}O$ | $C_{18}H_{29}NO_2$ | 292.2276 | - |

| | | | | |
|---|---|---|---|---|
| 4-Decanone | $C_{10}H_{20}O$ | $C_{18}H_{29}NO_2$ | 292.2276 | - |
| 3-Decanone | $C_{10}H_{20}O$ | $C_{18}H_{29}NO_2$ | 292.2276 | - |
| 2-Undecanone | $C_{11}H_{22}O$ | $C_{19}H_{31}NO_2$ | 306.2433 | - |
| *Aromatic ketones* | | | | |
| 1-Phenylethanone | $C_8H_8O$ | $C_{16}H_{17}NO_2$ | 256.1338 | - |
| 1-Phenylpropan-2-one | $C_9H_{10}O$ | $C_{17}H_{19}NO_2$ | 270.1494 | - |
| 1-(2-Methylphenyl)ethanone | $C_9H_{10}O$ | $C_{17}H_{19}NO_2$ | 270.1494 | - |
| 1-(4-Methylphenyl)ethanone | $C_9H_{10}O$ | $C_{17}H_{19}NO_2$ | 270.1494 | - |
| 4-Phenyl-3-buten-2-one | $C_{10}H_{10}O$ | $C_{18}H_{19}NO_2$ | 282.1494 | - |
| 1-Phenyl-2-butanone | $C_{10}H_{12}O$ | $C_{18}H_{21}NO_2$ | 284.1650 | - |
| 1-(3,4-Dimethyphenyl)-ethanone | $C_{10}H_{12}O$ | $C_{18}H_{21}NO_2$ | 284.1650 | - |
| *Cyclic ketones* | | | | |
| Cyclohexanone | $C_6H_{10}O$ | $C_{14}H_{19}NO_2$ | 234.1494 | - |

[a] Endogenous carbonyl species were compiled from previous studies of *S. cerevisiae* metabolism[8,9]

[b] The product of dopamine+2MB is isomeric with that of dopamine+3MB ($m/z$ + $H^+$ = 222.1494). The identity of the putative THIQ product was deduced to be dopamine+3MB since growth on L-leucine, rather than L-isoleucine, generates the corresponding LC-MS peak (refer to Supplementary Fig. 12). Further, α-substituted carbonyls, such as 2MB, are not well tolerated by NCS[10].

**Table S2 – QTOF-MS analysis of substituted tetrahydroisoquinolines synthesized from supplemented amino acids**

| THIQ product | THIQ formula | Retention time (min) | Expected THIQ mass (*m/z* + $H^+$) | Observed THIQ mass (*m/z* + $H^+$) | Mass error (ppm) |
|---|---|---|---|---|---|
| **25** | $C_{12}H_{17}NO_2S$ | 3.16 | 240.1058 | 240.1061 | 1.2 |
| **26** | $C_{14}H_{21}NO_2S$ | 5.13 | 268.1371 | 268.1370 | 0.4 |
| **28** | $C_{16}H_{17}NO_4$ | 3.17 | 288.1236 | 288.1230 | 2.1 |
| **10** | $C_{18}H_{21}NO_4$ | 4.81 | 316.1549 | 316.1542 | 2.2 |
| **31** | $C_{11}H_{15}NO_2$ | 1.98 | 194.1181 | 194.1185 | 2.1 |
| **32** | $C_{13}H_{19}NO_2$ | 4.64 | 222.1494 | 222.1492 | 0.9 |
| **35** | $C_{12}H_{17}NO_2$ | 2.91 | 208.1338 | 208.1332 | 2.9 |
| **36** | $C_{14}H_{21}NO_2$ | 5.06 | 236.1650 | 236.1645 | 2.1 |
| **39** | $C_{13}H_{19}NO_2$ | 4.63 | 222.1494 | 222.1499 | 2.2 |
| **40** | $C_{15}H_{23}NO_2$ | 5.35 | 250.1807 | 250.1803 | 1.6 |
| **43** | $C_{14}H_{21}NO_2$ | 5.34 | 236.1650 | 236.1645 | 2.1 |
| **44** | $C_{16}H_{25}NO_2$ | 5.52 | 264.1963 | 264.1958 | 1.9 |

**Table S3 – Plasmids utilized in this study**

| Plasmid | Description | Source or reference |
|---|---|---|
| pCAS-G418 | $P_{RNR2}$-*cas9*$_{NLS}$-$T_{CYC1}$, pUC, 2μ, $P_{tRNA\_Tyr}$-3′HDV-gRNA-Scaffold-$T_{SNR52}$, $P_{TEF1}$-*kanMX*-$T_{TEF1}$ | [11] |
| pBOT-His | CEN6/ARS4$^{ori}$, pMB1$^{ori}$, Amp$^{R}$, Kan$^{R}$, *HIS3*, $P_{TEF1}$-*GFP*$^{S65T}$-$T_{PGI1}$ | [12] |
| pBOT-NdNCS | CEN6/ARS4$^{ori}$, pMB1$^{ori}$, Amp$^{R}$, Kan$^{R}$, *HIS3*, $P_{TEF1}$-*NdNCS*-$T_{PGI1}$ | [12] |
| pBOT-ScNCS | CEN6/ARS4$^{ori}$, pMB1$^{ori}$, Amp$^{R}$, Kan$^{R}$, *HIS3*, $P_{TEF1}$-*ScNCS*-$T_{PGI1}$ | [12] |
| pCAS-Hyg | $P_{RNR2}$-*cas9*$_{NLS}$-$T_{CYC1}$, pUC, 2μ, $P_{tRNA\_Tyr}$-3′HDV-gRNA-Scaffold-$T_{SNR52}$, $P_{TEF1}$-*HphNTI*-$T_{TEF1}$ | This study |
| pJET-LP5.T3 | pMB1$^{ori}$, Amp$^{R}$, LP5.T3 | This study |
| pBSC009 | ColE1, Kan$^{R}$, *LEU2*, $P_{TDH3}$-*CjNCS*-$T_{ENO2}$ | This study |
| pPSG325 | ColE1, Kan$^{R}$, *LEU2*, $P_{TDH3}$-*CjNCSΔN*$_{20}$-$T_{ENO2}$ | This study |
| pBSC011 | ColE1, Kan$^{R}$, *LEU2*, $P_{TDH3}$-*CjNCSΔN*$_{35}$-$T_{ENO2}$ | This study |
| pPSG834 | CEN6/ARS4$^{ori}$, ColE1, Kan$^{R}$, *HIS3*, $P_{CCW12}$-*CYP76AD1*$^{W13L\ F309L}$-$T_{ENO2}$ | This study |
| pPSG835 | CEN6/ARS4$^{ori}$, ColE1, Kan$^{R}$, *HIS3*, $P_{CCW12}$-*CYP76AD5*-$T_{ENO2}$ | This study |
| pPSG836 | CEN6/ARS4$^{ori}$, ColE1, Kan$^{R}$, *HIS3*, $P_{CCW12}$-*CYP76AD6*-$T_{ENO2}$ | This study |
| pPSG450 | CEN6/ARS4$^{ori}$, ColE1, Kan$^{R}$, *HIS3*, $P_{TEF1}$-*EcCYP80B1*-$T_{TDH3}$, $P_{TDH3}$-*Ps6OMT*-$T_{ADH1}$, $P_{PGK1}$-*Ps4′OMT2*-$T_{ENO1}$, $P_{TEF2}$-*PsCNMT*-$T_{SSA1}$, $P_{HHF1}$-*AtATR2*-$T_{ENO2}$, | This study |

**Table S4 – Strains utilized in this study**

| ID | Description | Parent | Manipulation | Reference |
|---|---|---|---|---|
| BY4741 | Quadruple auxotroph | - | - | |
| 1373 | Dopamine producer | BY4741 | $P_{TDH3}$-*CYP76AD1*$^{W13L\ F309L}$-$T_{TDH1}$ $P_{CCW12}$-*PpDODC*-$T_{ADH1}$ $P_{PGK1}$-*ARO4*$^{FBR}$-$T_{PGK1}$ | [5] |
| LP165 | *Nd*NCS base strain | 1373 | $P_{TEF1}$-*NdNCS*-$T_{PGI1}$ | This study |
| LP167 | *Sc*NCS base strain | 1373 | $P_{TEF1}$-*ScNCS*-$T_{PGI1}$ | This study |
| LP172 | *Nd*NCSΔ$N_{20}$ base strain | LP165 | $P_{TEF1}$-*NdNCSΔN*$_{20}$-$T_{PGI1}$ | This study |
| LP173 | *Sc*NCSΔ$N_{20}$ base strain | LP167 | $P_{TEF1}$-*ScNCSΔN*$_{20}$-$T_{PGI1}$ | This study |
| LP273 | *Nd*NCS-GFP fusion | 1373 | $P_{TEF1}$-*NdNCS-GFP*-$T_{PGI1}$ | This study |
| LP274 | *Nd*NCSΔ$N_{20}$-GFP fusion | 1373 | $P_{TEF1}$-*NdNCSΔN*$_{20}$-*GFP*-$T_{PGI1}$ | This study |
| LP275 | *Sc*NCS-GFP fusion | 1373 | $P_{TEF1}$-*ScNCS-GFP*-$T_{PGI1}$ | This study |
| LP276 | *Sc*NCSΔ$N_{20}$-GFP fusion | 1373 | $P_{TEF1}$-*ScNCSΔN*$_{20}$-*GFP*-$T_{PGI1}$ | This study |
| LP277 | GFP control | 1373 | $P_{TEF1}$-*GFP*-$T_{PGI1}$ | This study |
| LP383 | *Cj*NCS base strain | 1373 | $P_{TEF1}$-*CjNCS*-$T_{PGI1}$ | This study |
| LP384 | *Cj*NCSΔ$N_{20}$ base strain | 1373 | $P_{TEF1}$-*CjNCSΔN*$_{20}$-$T_{PGI1}$ | This study |
| LP385 | *Cj*NCSΔ$N_{35}$ base strain | 1373 | $P_{TEF1}$-*CjNCSΔN*$_{35}$-$T_{PGI1}$ | This study |
| LP99 | *ald2*Δ *ald3*Δ mutant | LP165 | *ald2*Δ *ald3*Δ | This study |
| LP101 | *ald4*Δ mutant | LP165 | *ald4*Δ | This study |
| LP102 | *ald5*Δ mutant | LP165 | *ald5*Δ | This study |
| LP103 | *ald6*Δ mutant | LP165 | *ald6*Δ | This study |
| LP109 | *adh1*Δ mutant | LP165 | *adh1*Δ | This study |
| LP100 | *adh2*Δ mutant | LP165 | *adh2*Δ | This study |
| LP104 | *adh3*Δ mutant | LP165 | *adh3*Δ | This study |
| LP105 | *adh4*Δ mutant | LP165 | *adh4*Δ | This study |
| LP106 | *adh5*Δ mutant | LP165 | *adh5*Δ | This study |
| LP107 | *adh6*Δ mutant | LP165 | *adh6*Δ | This study |
| LP121 | *adh7*Δ mutant | LP165 | *adh7*Δ | This study |
| LP108 | *sfa1*Δ mutant | LP165 | *sfa1*Δ | This study |
| LP225 | *ari1*Δ mutant | LP165 | *ari1*Δ | This study |
| LP226 | *ygl039w*Δ mutant | LP165 | *ygl039w*Δ | This study |
| LP227 | *ydr541c*Δ mutant | LP165 | *ydr541c*Δ | This study |
| LP228 | *gre2*Δ mutant | LP165 | *gre2*Δ | This study |
| LP229 | *ypr1*Δ mutant | LP165 | *ypr1*Δ | This study |
| LP230 | *gcy1*Δ mutant | LP165 | *gcy1*Δ | This study |
| LP231 | *aad14*Δ mutant | LP165 | *aad14*Δ | This study |
| LP232 | *aad3*Δ mutant | LP165 | *aad3*Δ | This study |
| LP192 | *adh1*Δ-*noxE* mutant | LP109 | $P_{TEF1}$-*LlnoxE*-$T_{IDP1}$ | This study |
| LP295 | (*S*)-Norcoclaurine strain | LP225 | *ald4*Δ | This study |
| LP296 | (*S*)-Norcoclaurine strain | LP225 | *ald6*Δ | This study |
| LP297 | (*S*)-Norcoclaurine strain | LP225 | *ygl039w*Δ | This study |
| LP298 | (*S*)-Norcoclaurine strain | LP225 | *ydr541c*Δ | This study |
| LP299 | (*S*)-Norcoclaurine strain | LP225 | *adh6*Δ | This study |
| LP300 | (*S*)-Norcoclaurine strain | LP225 | *ypr1*Δ | This study |

| | | | | |
|---|---|---|---|---|
| LP316 | (*S*)-Norcoclaurine strain | LP295 | *adh6*Δ | This study |
| LP317 | (*S*)-Norcoclaurine strain | LP295 | *ypr1*Δ | This study |
| LP318 | (*S*)-Norcoclaurine strain | LP295 | *ydr541c*Δ | This study |
| LP328 | (*S*)-Norcoclaurine strain | LP316 | *NdNCS*::$NdNCS\Delta N_{20}$ | This study |
| LP348 | (*S*)-Norcoclaurine strain | LP328 | 3×$NdNCS\Delta N_{20}$ (4× total) | This study |
| LP355 | (*S*)-Norcoclaurine strain | LP348 | *ypr1*Δ | This study |
| LP357 | (*S*)-Norcoclaurine strain | LP355 | *adh3*Δ | This study |
| LP358 | (*S*)-Norcoclaurine strain | LP355 | *ydr541c*Δ | This study |
| LP359 | (*S*)-Norcoclaurine strain | LP355 | *adh2*Δ | This study |
| LP360 | (*S*)-Norcoclaurine strain | LP355 | *adh5*Δ | This study |
| LP361 | (*S*)-Norcoclaurine strain | LP355 | *adh7*Δ | This study |
| LP362 | (*S*)-Norcoclaurine strain | LP355 | *sfa1*Δ | This study |
| LP363 | (*S*)-Norcoclaurine strain | LP355 | *aad3*Δ | This study |
| LP364 | (*S*)-Norcoclaurine strain | LP355 | *aad14*Δ | This study |
| LP369 | (*S*)-Norcoclaurine strain | LP358 | 2×$NdNCS\Delta N_{20}$ (6× total) | This study |
| LP374 | (*S*)-Norcoclaurine strain | LP369 | $P_{TDH3}$-*TYR1*-$T_{TDH1}$ | This study |
| LP375 | (*S*)-Norcoclaurine strain | LP374 | $P_{TDH3}$-$ARO7^{FBR}$-$T_{TDH1}$ | This study |
| LP376 | (*S*)-Norcoclaurine strain | LP375 | $P_{TDH3}$-*ARO10*-$T_{TDH1}$ | This study |
| LP377 | (*S*)-Norcoclaurine strain | LP376 | *pha2*Δ | This study |
| LP379 | (*S*)-Norcoclaurine strain | LP377 | *trp3*Δ | This study |
| LP381 | (*S*)-Norcoclaurine strain | LP379 | 2×$NdNCS\Delta N_{20}$ (8× total) | This study |
| LP382 | (*S*)-Norcoclaurine strain | LP381 | $P_{TDH3}$-*ARO2*-$T_{TDH1}$ | This study |
| LP386 | (*S*)-Norcoclaurine strain | LP382 | *aad3*Δ | This study |
| LP412 | (*S*)-Norcoclaurine strain | LP386 | $P_{TDH3}$-$CjNCS\Delta N_{35}$-$T_{TDH1}$ | This study |
| LP430 | (*S*)-Norcoclaurine strain | LP412 | Deletion of 7×$NdNCS\Delta N_{20}$ | This study |
| LP442 | (*S*)-Norcoclaurine strain | LP430 | Deletion of final 1×$NdNCS\Delta N_{20}$ | This study |
| LP456 | (*S*)-Norcoclaurine strain | LP442 | $P_{ALD4}$-*ALD4*-$T_{ALD4}$ | This study |
| LP474 | (*S*)-Norcoclaurine strain | LP456 | $P_{TEF1}$-$CjNCS\Delta N_{35}$-$T_{TDH1}$ | This study |
| LP476 | (*S*)-Norcoclaurine strain | LP474 | $P_{TEF1}$-$CYP76AD1^{W13L\ F309L}$-$T_{TDH1}$ | This study |
| LP477 | (*S*)-Norcoclaurine strain | LP474 | $P_{TDH3}$-$CYP76AD1^{W13L\ F309L}$-$T_{TDH1}$ | This study |
| LP478 | (*S*)-Norcoclaurine strain | LP474 | $P_{TEF1}$-*CYP76AD5*-$T_{TDH1}$ | This study |
| LP479 | (*S*)-Norcoclaurine strain | LP474 | $P_{TDH3}$-*CYP76AD5*-$T_{TDH1}$ | This study |
| LP480 | (*S*)-Norcoclaurine strain | LP474 | $P_{TEF1}$-*CYP76AD6*-$T_{TDH1}$ | This study |
| LP481 | (*S*)-Norcoclaurine strain | LP474 | $P_{TDH3}$-*CYP76AD6*-$T_{TDH1}$ | This study |
| LP490 | (*S*)-Reticuline strain | LP478 | $P_{TEF1}$-*EcCYP80B1*-$T_{TDH3}$<br>$P_{TDH3}$-*Ps6OMT*-$T_{ADH1}$<br>$P_{PGK1}$-*Ps4′OMT2*-$T_{ENO1}$<br>$P_{TEF2}$-*PsCNMT*-$T_{SSA1}$<br>$P_{HHF1}$-*AtATR2*-$T_{ENO2}$ | This study |
| LP491 | (*S*)-Reticuline strain | LP490 | $P_{TDH3}$-*Ps4′OMT2*-$T_{TDH1}$ (2$^{nd}$ copy) | This study |
| LP492 | (*S*)-Reticuline strain | LP491 | Repair of BY4741 *SAL1* frameshift (G403L) | This study |
| LP494 | (*S*)-Reticuline strain | LP492 | *gre2*Δ | This study |
| LP495 | (*S*)-Reticuline strain | LP492 | *ald2*Δ *ald3*Δ | This study |
| LP498 | (*S*)-Reticuline strain | LP494 | *hfd1*Δ | This study |

| | | | | |
|---|---|---|---|---|
| LP499 | (*S*)-Reticuline strain | LP494 | $P_{TDH3}$-*Ps6OMT*-$T_{TDH1}$ (2$^{nd}$ copy) | This study |
| LP501 | (*S*)-Reticuline strain | LP499 | *hfd1*Δ | This study |

[a] Intermediate strains used in the construction of the final (*S*)-reticuline production host (LP501) are shaded.

**Table S5 – Expression cassettes utilized in this study**

| **Description** | **Cassette** | **Locus** |
|---|---|---|
| Testing *Nd*NCS | FgF20-$P_{TEF1}$-*NdNCS*-$T_{PGI1}$-FgF20 | FgF20 |
| Testing *Sc*NCS | FgF20-$P_{TEF1}$-*ScNCS*-$T_{PGI1}$-FgF20 | FgF20 |
| Testing *Nd*NCS$\Delta N_{20}$ | FgF20-$P_{TEF1}$-*NdNCS*$\Delta N_{20}$-$T_{PGI1}$-FgF20 | FgF20 |
| Testing *Sc*NCS$\Delta N_{20}$ | FgF20-$P_{TEF1}$-*ScNCS*$\Delta N_{20}$-$T_{PGI1}$-FgF20 | FgF20 |
| Testing *Cj*NCS | FgF20-$P_{TEF1}$-*CjNCS*-$T_{PGI1}$-FgF20 | FgF20 |
| Testing *Cj*NCS$\Delta N_{20}$ | FgF20-$P_{TEF1}$-*CjNCS*$\Delta N_{20}$-$T_{PGI1}$-FgF20 | FgF20 |
| Testing *Cj*NCS$\Delta N_{35}$ | FgF20-$P_{TEF1}$-*CjNCS*$\Delta N_{35}$-$T_{PGI1}$-FgF20 | FgF20 |
| *Nd*NCS-GFP fusion | FgF20-$P_{TEF1}$-*NdNCS-GFP*-$T_{PGI1}$-FgF20 | FgF20 |
| *Sc*NCS-GFP fusion | FgF20-$P_{TEF1}$-*ScNCS-GFP*-$T_{PGI1}$-FgF20 | FgF20 |
| *Nd*NCS$\Delta N_{20}$-GFP fusion | FgF20-$P_{TEF1}$-*NdNCS*$\Delta N_{20}$-*GFP*-$T_{PGI1}$-FgF20 | FgF20 |
| *Sc*NCS$\Delta N_{20}$-GFP fusion | FgF20-$P_{TEF1}$-*ScNCS*$\Delta N_{20}$-*GFP*-$T_{PGI1}$-FgF20 | FgF20 |
| GFP control | FgF20-$P_{TEF1}$-*GFP*-$T_{PGI1}$-FgF20 | FgF20 |
| Expression of *noxE* in *adh1*Δ | FgF7-$P_{TEF1}$-*LlnoxE*-$T_{IDP1}$-FgF7 | FgF7 |
| Expression of *NdNCS* or *NdNCS*$\Delta N_{20}$ | FgF20-LV3-$P_{TEF1}$-*NdNCS*-$T_{PGI1}$-LV5-FgF20<br>FgF20-LV3-$P_{TEF1}$-*NdNCS*$\Delta N_{20}$-$T_{PGI1}$-LV5-FgF20 | FgF20 |
| Overexpression of *TYR1* | USERXII-2-LV3-$P_{TDH3}$-*TYR1*-$T_{TDH1}$-LV5-USERXII-2 | USERXII-2 |
| Expression of *ARO7*$^{FBR}$ | FgF16-LV3-$P_{TDH3}$-*ARO7*$^{FBR}$-$T_{TDH1}$-LV5-FgF16 | FgF16 |
| Overexpression of *ARO10* | FgF18-LV3-$P_{TDH3}$-*ARO10*-$T_{TDH1}$-LV5-FgF18 | FgF18 |
| Overexpression of *ARO2* | FgF19-LV3-$P_{TDH3}$-*ARO2*-$T_{TDH1}$-LV5-FgF19 | FgF19 |
| Expression of 1$^{st}$ copy of *CjNCS*$\Delta N_{35}$ | FgF24-LV3-$P_{TDH3}$-*CjNCS*$\Delta N_{35}$-$T_{TDH1}$-LV5-FgF24 | FgF24 (*PDC6*) |
| Reintroduction of *ALD4* | 308a-LV3-$P_{ALD4}$-*ALD4*-$T_{ALD4}$-LV5-308a | 308a |
| Expression of 2$^{nd}$ copy of *CjNCS*$\Delta N_{35}$ | USERXII-5-LV3-$P_{TEF1}$-*CjNCS*$\Delta N_{35}$-$T_{PGI1}$-LV5-USERXII-5 | USERXII-5 |
| Expression of *CYP76AD5* | 1309a-LV3-$P_{TEF1}$-*CYP76AD5*-$T_{TDH1}$-LV5-1309a | 1309a |
| Introduction of reticuline biosynthesis pathway | 106a-LV3-$P_{TEF1}$-*EcNMCH*-$T_{TDH3}$-$P_{TDH3}$-*Ps6OMT*-$T_{ADH1}$-$P_{PGK1}$-*Ps4′OMT2*-$T_{ENO1}$-$P_{TEF2}$-*PsCNMT*-$T_{SSA1}$-$P_{HHF1}$-*AtATR2*-$T_{ENO2}$-LV5-106a | 106a |
| Expression of 2$^{nd}$ copy of *Ps4′OMT* | 416d-LV3-$P_{TDH3}$-*Ps4′OMT2*-$T_{TDH1}$-LV5-416d | 416d |
| Expression of 2$^{nd}$ copy of *Ps6OMT* | 511b-LV3-$P_{TDH3}$-*Ps6OMT*-$T_{TDH1}$-LV5-511b | 511b |

**Table S6 – *S. cerevisiae* integration sites utilized in this study**

| Target site ID | Target site sequence[a] | Reference |
|---|---|---|
| FgF7 | TATCCTGAATGTTCTCTCCCAGG | [12,13] |
| FgF16 | TGTACCAAAAGTTATCCTGTAGG | [12,13] |
| FgF18 | ATAGAATTACTATTGAAGAGTGG | [12,13] |
| FgF19 | ATTCACTCTGCTAAGATTATCGG | [12,13] |
| FgF20 | GTTAGAGCTGTTACAAGTTACGG | [12,13] |
| FgF24 (*PDC6*) | GTACAACGAAATCCAGACCTGGG | [12,13] |
| USERXII-2 | TCGAGAGAGTCGCCGATAGTAGG | [14] |
| USERXII-5 | TTGTCACAGTGTCACATCAGCGG | [14] |
| 106a | ATACGGTCAGGGTAGCGCCCTGG | [4] |
| 308a | CACTTGTCAAACAGAATATAAGG | [4] |
| 416d | TAGTGCACTTACCCCACGTTCGG | [4] |
| 511b | CAGTGTATGCCAGTCAGCCACGG | [4] |
| 1309a | CCTGTGGTGACTACGTATCCAGG | [4] |
| *AAD3* | CAGGCGGAATGTAATAGGTGCGG | This study |
| *AAD14* | TCGTATTCAAGGAAACCGGGGGG | This study |
| *ALD2+ALD3* | GCTCAAGAATGTTCATATAAAGG | This study |
| *ALD4* | GGGTGTAGGTAAGCAGAATGAGG | This study |
| *ALD5* | TTAGAGTTTTTCGATGAGAATGG | This study |
| *ALD6* | ACACCGTTCGAGGTCAAGCCTGG | This study |
| *ADH1* | CTCTAATGAGCAACGGTATACGG | This study |
| *ADH2* | GCACTCTATTTATATGTGATAGG | This study |
| *ADH3* | AGCGAGTGTTCCTTTCTAAAAGG | This study |
| *ADH4* | CAATTGCTTTGTAGAGTTAACGG | This study |
| *ADH5* | TTACAATCTAGACAATACGAAGG | This study |
| *ADH6* | CACTTCACCTCGAGAACTGTTGG | This study |
| *ADH7* | AATCCCAATGTCATTTAATGCGG | This study |
| *ARI1* | AATTAGCATAGGATTTTCCGCGG | This study |
| *GCY1* | TAGGGAATTAAGGAGAGCAGCGG | This study |
| *GRE2* | TAGAATACGGAATTTTCTCGCGG | This study |
| *HFD1* | AGGCATATTGATTATCTAAAAGG | This study |
| *NdNCS* (N-terminus) | TTGGGTTGTGAAATTTCCCAAGG | This study |
| *NdNCS* ($T_{PGI1}$-LP5) | AGTTAGGTCTGGTATACTGGAGG | This study |
| *PHA2* | TCAGCGACAAAAGTAAACAGTGG | This study |
| *SFA1* | GTAATAATGGAATTTCATAGAGG | This study |
| T3 | GCCAGTCAGAACACTAGAGGCGG | This study |
| *TRP3* | TCTATCGGGAATTACCACCAGGG | This study |
| *YDR541C* | GGCACCCAAAATAGATAGAGTGG | This study |
| *YGL039W* | GGGTTTGGCACAATTTGGCTTGG | This study |

| | | |
|---|---|---|
| *YPR1* | GAAACCCAACACTGGAATGGAGG | This study |

[a] PAMs are underlined.

#### Table S7 – Genes and synthetic DNAs synthesized in this study

| DNA | Source | Ref. | Codon opt. | Sequence (5′ – 3′)[a] |
|---|---|---|---|---|
| *CYP76AD1** | *Beta vulgaris* | [5] | Yes | ATGGATCACGCAACATTAGCTATGATATTAGCAATTTTGTTTATTTCATTTCATTTTATT<br>AAATTATTATTCTCACAACAAACAACAAAATTGTTACCTCCAGGTCCTAAACCTTTACC<br>AATTATTGGTAATATCTTGGAAGTAGGAAAGAAGCCTCATAGATCATTTGCAAATTTGG<br>CTAAGATCCACGGTCCTTTAATCTCTTTGAGGTTAGGATCAGTTACCACCATTGTTGTAT<br>CATCTGCTGATGTCGCCAAAGAGATGTTCTTGAAGAAGGATCACCCTTTATCAAACAGG<br>ACAATTCCAAATTCAGTAACTGCTGGTGACCACCACAAATTGACTATGTCTTGGTTGCC<br>TGTTTCACCTAAATGGAGAAACTTCAGAAAGATAACTGCTGTTCATTTATTGTCTCCAC<br>AAAGATTAGATGCATGCCAGACCTTTAGACATGCAAAGGTTCAGCAATTATATGAATA<br>CGTTCAAGAGTGCGCTCAGAAGGGTCAGGCAGTTGACATTGGTAAAGCTGCCTTCACA<br>ACTTCTTTGAATTTATTGTCTAAATTATTCTTTTCTGTTGAATTGGCCCATCATAAATCA<br>CATACCTCTCAAGAGTTTAAGGAATTAATTTGGAATATTATGGAAGACATCGGTAAACC<br>AAACTACGCCGATTACTTCCCAATTTTAGGTTGCGTCGATCCATCAGGAATTAGAAGAA<br>GGTTGGCATGTTCTTTTGATAAGTTAATTGCAGTATTTCAAGGAATAATATGTGAGAGG<br>TTAGCTCCAGATTCATCAACTACAACTACTACAACTACAGATGATGTCTTGGACGTATT<br>ATTGCAATTGTTTAAGCAGAATGAATTAACTATGGGAGAGATTAACCACTTGTTAGTCG<br>ATATTTTCGATGCCGGTACTGATACTACATCATCTACCTTAGAGTGGGTTATGACTGAG<br>TTAATAAGAAACCCTGAAATGATGGAAAAGGCCCAAGAGGAAATAAAACAGGTATTG<br>GGTAAGGATAAGCAAATCCAAGAGTCAGACATAATAAATTTGCCATACTTGCAAGCAA<br>TCATTAAGGAAACTTTGAGGTTGCACCCACCTACAGTTTTCTTGTTGCCTAGAAAAGCC<br>GACACAGACGTAGAATTGTACGGATACATTGTTCCAAAGGATGCTCAAATTTTGGTAA<br>ACTTGTGGGCCATAGGTAGAGATCCAAATGCTTGGCAAAATGCCGATATTTTCTCACCT<br>GAAAGATTTATAGGTTGTGAGATTGATGTAAAGGGAAGAGATTTTGGTTTATTGCCTTT<br>TGGTGCTGGTAGAAGAATTTGCCCTGGTATGAATTTGGCCATAAGGATGTTGACATTGA<br>TGTTGGCCACATTATTGCAATTCTTCAACTGGAAATTAGAGGGAGACATTTCACCTAAA<br>GATTTGGACATGGATGAAAAGTTTGGTATAGCCTTGCAGAAAACTAAACCTTTAAAATT<br>GATCCCAATTCCAAGATACGGATCCTAA |
| *CYP76AD5* | *Beta vulgaris* | [2] | Yes | ATGGATAATACAACATTAGCCTTAATTTTATCATCATTGTTCGTATGTTTCCAATTGATA<br>AGGTCATTTATTAATCATGCTAAGAAATCAAATAAATTGCCACCTGGTCCTAAGAGAAT<br>GCCAATTTTCGGTAATATCTTTGACTTGGGAGAGAAGCCTCATAGATCATTCGCAAATT<br>TGGCCAAAATTCATGGTCCTTTGGTTTCATTGCAATTGGGTTCTGTTACAACCGTAGTAG<br>TATCATCTGCCGATGTTGCAAAGGAAATGTTCTTAAAGAACGACCAAGCTTTGGCTAAT<br>AGGACTATACCAGACTCTGTTAGAGCTGGTGATCACGATAAATTATCAATGTCATGGTT<br>ACCAGTATCAGCCAAGTGGAGGAATTTGAGGAAAATATCTGCTGTACAGTTGTTATCA<br>ACCCAGAGATTGGATGCATCTCAAGCTCATAGGCAGTCTAAAGTACAACAATTGTTGG<br>AGTATGTTCACGATTGCTCAAAGAAAGGTCAACCAGTTGACATTGGAAGAGCCGCTTTT |

| | | | | |
|---|---|---|---|---|
| | | | | ACTACTTCTTTGAATTTATTGTCTAACACTTTCTTCTCTGTAGAATTGGCCTCACACGAG<br>TCTTCAGCATCACAAGAATTTAAACAGTTAATGTGGAACATTATGGAAGAAATCGGTA<br>GACCAAATTATGCTGACTTCTTTCCTATATTGGGATATTTGGACCCATTTGGTATTAGAA<br>GAAGATTAGCAGGTTACTTCGATCAATTAATTGCCGTCTTTCAAGATATTATAGGTGAG<br>AGACAGAAGATAAGATCCGCCAACTTGTCTGGAGGTAAACAAACAACAAACGATATCT<br>TAGACACTTTGTTAAACTTGTACGATGAAAAGGAGTTGTCTATGGGCGAGATTAACCAC<br>TTATTGGTAGATATATTTGATGCCGGAACTGATACCACTGCATCAACCTTGGAATGGGC<br>AATGGCCGAATTGGTTAAGAACCCTGATATGATGGTAAAGGTACAGGATGAGATTGAA<br>CAAGCAATAGGAAAGGGATGTTCTATGGTACAAGAGTCAGACATCTCTAAATTACCTT<br>ATTTACAAGCCATTATTAAGGAAACTTTGAGATTACACCCTCCTACAGTCTTCTTGTTGC<br>CAAGAAAGGCCGATGCCGATGTAGAGTTATACGGTTATGTAGTCCCCAAGAATGCTCA<br>AGTCTTGGTTAATTTGTGGGCTATCGGAAGAGATCCTAAGGTCTGGAAGAATCCAGAG<br>GTTTTCTCACCAGAAAGATTCTTGGAAAGTAACATTGATTACAAAGGTAGGGACTTCGA<br>GTTGTTGCCTTTTGGTGCTGGTAGGAGAATATGTCCAGGTTTGACCTTAGCTTACAGAA<br>TGTTGAACCTGATGATGGCTAACTTCTTGCATTCTTATGACTGGAAGTTAGAAGATGGT<br>ATGCATCCAAAAGATTTGGACATGGACGAGAAGTTTGGTATCACTTTGCAGAAAGTCA<br>AGCCATTGCAGGTCATCCCAGTTCCAAGGAAAGGATCCTAA |
| *CYP76AD6* | *Beta vulgaris* | 2 | Yes | ATGGATAACGCAACACTTGCTGTGATCCTTTCCATTTTGTTTGTGTTTTACCACATTTTC<br>AAATCCTTTTTCACCAATTCTTCATCTCGTAGGCTTCCTCCTGGTCCCAAACCCGTGCCA<br>ATTTTTGGCAACATTTTCGATCTTGGCGAAAAGCCTCATCGATCTTTTGCCAATCTATCT<br>AAAATTCACGGCCCTTTGATTAGCCTAAAGTTAGGAAGTGTAACAACTATTGTTGTTTC<br>CTCGGCCTCTGTGGCCGAGGAAATGTTCCTTAAAAATGACCAAGCACTTGCTAACCGAA<br>CCATTCCTGACTCGGTTAGGGCTGGTGACCACGACAAATTATCCATGTCGTGGTTGCCT<br>GTTTCCCAAAAATGGAGAAATATGAGAAAAATCTCCGCTGTCCAATTACTCTCCAACCA<br>AAAACTTGATGCTAGTCAACCTCTTAGACAAGCTAAGGTGAAACAACTTTTATCATACG<br>TACAAGTTTGTTCCGAAAAAATGCAACCCGTCGATATTGGACGGGCCGCATTTACAACG<br>TCACTTAATTTATTATCAAACACATTTTTCTCAATCGAATTAGCAAGTCATGAATCTAGT<br>GCTTCCCAAGAGTTTAAACAACTCATGTGGAATATTATGGAGGAAATTGGAAGGCCTA<br>ATTATGCTGATTTTTTCCCTATTCTTGGTTACATTGATCCCTTTGGTATAAGACGTCGTTT<br>GGCTGGTTACTTTGATAAACTCATTGATGTTTTCCAAGACATTATTCGTGAAAGACAAA<br>AGCTTCGATCTTCTAATTCTTCCGGCGCAAAACAAACAAATGACATTCTTGATACTCTT<br>CTTAAACTCCATGAAGATAATGAGTTGAGTATGCCTGAAATTAATCACCTTCTCGTGGA<br>TATCTTTGACGCCGGAACAGACACAACAGCAAGCACATTAGAATGGGCGATGGCCGAA<br>CTTGTGAAAAACCCGGAAATGATGACTAAAGTTCAAATTGAAATCGAACAAGCTCTTG<br>GAAAAGATTGCTTAGACATACAAGAATCCGACATCTCAAAACTACCTTATTTACAAGCC<br>ATTATAAAAGAAACGTTACGTTTACACCCTCCTACTGTGTTTTTGCTGCCTCGAAAGGC<br>AGACAATGACGTAGAGTTATATGGCTACGTTGTACCAAAGAATGCTCAAGTCCTTGTCA<br>ATCTTTGGGCAATTGGTCGTGATCCAAAGGTATGGAAAAATCCGGAAGTATTTTCTCCT<br>GAAAGGTTTTTAGATTGCAATATCGATTATAAAGGACGAGATTTCGAACTTTTACCCTT<br>TGGTGCTGGTAGAAGGATATGCCCTGGACTTACTTTGGCATATAGAATGTTGAACTTGA |

| | | | | |
|---|---|---|---|---|
| | | | | TGTTGGCTACTCTTCTTCAAAACTACAATTGGAAACTTGAAGATGGTATCAATCCTAAG<br>GATTTAGACATGGATGAGAAATTTGGGATTACATTGCAAAAGGTTAAACCTCTTCAAGT<br>TATTCCAGTTCCCAGAAACGGATCCTAA |
| *NdNCS* | *Nandina domestica* | [12] | Yes | ATG**AGATCTGGTATTGTCTTCTTGGTCTTGTTCTTCTTGGGTTGTGAAATTTCCCA**<br>**AGGT**AGACAATTGTTAGAGTCTAGATTATTCAGAAAGTCCACCATTAGAAAGGTCTTA<br>CACCACGAATTATCCGTTGCCGCTTCCGCTCAAGAAGTTTGGGATGTTTACTCCTCTCCA<br>GAATTGCCAAAGCATTTGCCAGAGATTTTACCAGGTGCTTTCAAAAAAGTCGTCGTTAC<br>TGGTGACGGTGGTGTTGGTACTGTTATTGAGATGATTTTCCCACCAGGTGTCGTCCCAC<br>ACAGATACAAAGAAAAATTCGTTTTAATTGATGACGAAAAATTTTTGAAGAAGGTTGA<br>AATGATCGAAGGTGGTTACTTGGATATGGGTTGTACTTTTTACATGGACACTATTCAAA<br>TCATTCCAACCGGTCCTGATTCTTGCATTATCAAGTCTTCTACTGAATACTACGTCAAGC<br>CAGAGTTCGCTGATAAGGTTGTCCCATTAATTTCTACTGTTCCATTACAAGCTATGGCTG<br>AAGCTATCGCTAAGATCGTTTTGGAAAACAAGGCTAAGCATAAGGGTTTTATCGAAAT<br>CGGCTGA |
| *ScNCS* | *Sanguinaria canadensis* | [12] | Yes | ATG**AGATCTGGTATTGTCTTTTTGGTTTTGTTCTTCTTAGGTTGTGAAATTTCCCAA**<br>**GGT**AGACAATTGTTGGAATCTAGATTGTTCAGAAAGTCTACTATTCAAAAGGTTTTGCA<br>CCACGAATTGCCAGTCGCTGCTTCTGCTCAAGAAGTTTGGGATGTCTATTCTTCCCCAG<br>AATTGCCAAAACACTTGCCAGAAATTTTGCCAGGTGCCTTCGAAAAGGTTGTCGTCACC<br>GGTGATGGTGGTGTTGGTACTGTTTTGGAAATGGTTTTTCCACCAGGTGAAGTTCCAAG<br>ATCCTACAAAGAAAAGTTCGTCTTAATTGACGATGAGCAATTGTTGAAGAAGGTTGAA<br>ATGATCGAAGGTGGTTACTTGGATATGGGTTGTACTTTCTACATGGACACTATCCAAAT<br>TGTCCCAACCGGTCCAGACTCCTGTATCATCAAGTCTTCTACTGAATACTATGTCAAGC<br>CTGAATTCGCTGATAAGGTTGTTCCATTGATTTCTACTATCCCATTGCAAGCTATGGCCG<br>AGGCCATCTCTAACATTGTCTTAGCTAACAAGGCCAAGAACAAGTCTATTATTATCGAA<br>ATTGGCTGA |
| *CjNCS* | *Coptis japonica* | This study | Yes | ATG**AGGATGGAAGTTGTATTGGTCGTCTTCTTGATGTTTATTGGTACTATTAACTG**<br>**TGAGAGGTTGATTTTCAATGGTAGACCATTGTTGCACAGAGTCACAAAA**GAAGAA<br>ACTGTAATGTTATATCACGAATTGGAAGTCGCAGCCTCAGCTGACGAAGTATGGTCTGT<br>CGAAGGTTCTCCTGAATTAGGTTTACATTTACCAGATTTGTTACCAGCTGGAATATTCG<br>CAAAGTTTGAGATAACTGGAGATGGTGGAGAAGGTTCTATTTTGGACATGACATTTCCA<br>CCTGGTCAATTCCCTCATCATTATAGAGAGAAGTTCGTTTTCTTCGACCATAAGAATAG<br>ATATAAGTTGGTTGAACAAATTGACGGTGATTTCTTTGATTTGGGAGTAACCTACTACA<br>TGGACACTATTAGGGTCGTCGCAACAGGTCCTGATTCTTGTGTCATCAAATCAACAACT<br>GAATATCATGTTAAACCAGAATTTGCTAAGATTGTTAAACCTTTAATCGATACAGTTCC<br>ATTAGCCATTATGTCAGAGGCAATAGCTAAGGTAGTATTGGAAAATAAGCACAAATCT<br>TCTGAGTAA |
| LP5.T3 | Random DNA sequence | This study | No | TGACTACCTGTTACTAAGCACATCGTGGTTTTCAGATATCGTGCACGTAGGGTTGCACC<br>GCACGCATGTGGAATTAGTGGCGAAGTACGATTCCACGACCGACGTACGATTCAACTA<br>TGCGGACGTGACGAGCTTCTTTTATATGCTTCGCCCGCCGGACCGGCCTCGTGATGGGG<br>TAGCTGCGCATAAGCTTATGACAATTAACGAGTGTGTACTCGTTTTATCATCTCACAGT |

| | |
|---|---|
| | TAAAGTCGGGAGAATAGGAGCCGCTACACACGAGTTACCACATCTGCCAGTCAGAACACTAGAGGcggAGACCTAACTGAGATACTGCCATAGACGACTACCCATCCCTCTGGGCCTTAGATAGCCGGATACAGTGACTTTGAAAGGTTTGTGGGGTACAGCTATGACTTGCTTAGCTGCGTGTGGGGGAAGGAACTTTTGCGTGTTAGTATGTTGACCCGTGTATTACGCATGCGGGTAGATTATGTAGGTAGAGACATCCAGGTCAAGTTCTCGACCTTCTCGTGGGAGGTGAACCAGTTCACTATAGGACCATTCCGTTCGAGCATGGCACTAAGTACGACCTCC |

[a] NCS regions corresponding to $\Delta N_{20}$ truncations are bolded and underlined and regions corresponding to $\Delta N_{35}$ truncations are bolded. The synthetic T3 Cas9 target site is underlined in LP5.T3.
